# Members of a highly widespread bacteriophage family are hallmarks of metabolic syndrome gut microbiomes

**DOI:** 10.1101/2021.03.30.437683

**Authors:** Patrick A. de Jonge, Koen Wortelboer, Torsten P.M. Scheithauer, Bert-Jan H. van den Born, Aeilko H. Zwinderman, Franklin L. Nobrega, Bas E. Dutilh, Max Nieuwdorp, Hilde Herrema

## Abstract

There is significant interest in altering the course of cardiometabolic disease development via the gut microbiome. Nevertheless, the highly abundant phage members -which impact gut bacteria- of the complex gut ecosystem remain understudied. Here, we characterized gut phageome changes associated with metabolic syndrome (MetS), a highly prevalent clinical condition preceding cardiometabolic disease. MetS gut phageome populations exhibited decreased richness and diversity, but larger inter-individual variation. These populations were enriched in phages infecting *Bacteroidaceae* and depleted in those infecting *Ruminococcaeae*. Differential abundance analysis identified eighteen viral clusters (VCs) as significantly associated with either MetS or healthy phageomes. Among these are a MetS-associated *Roseburia* VC that is related to healthy control-associated *Faecalibacterium* and *Oscillibacter* VCs. Further analysis of these VCs revealed the *Candidatus Heliusviridae*, a highly widespread gut phage lineage found in 90+% of the participants. The identification of the temperate *Ca. Heliusviridae* provides a novel starting point to a better understanding of the effect that phages have on their bacterial hosts and the role that this plays in MetS.

## Introduction

The human gut microbiome influences many (metabolic) processes, including digestion, the immune system^1^, and endocrine functions^2^. It is also involved in diseases such as type 2 diabetes^3^, fatty liver disease^4^ and inflammatory bowel disease^5^. Though studies of these gut microbiome effects on health and disease mostly focus on bacteria, increasing attention is devoted to bacteriophages (or phages).

Phages are viruses that infect bacteria. By infecting bacteria, they can significantly alter gut bacterial communities, mainly by integrating into bacterial genomes as prophages (lysogeny) or killing bacteria (lysis). Such alterations to bacterial communities in turn affect the interactions between bacteria and host, making phages part of an interactive network with bacteria and hosts. For example, an increase in phage lytic action is linked to decreased bacterial diversity in inflammatory bowel disease^6,7^, prophage integration into *Bacteroides vulgatus* modifies bacterial bile acid metabolism^8^, and dietary fructose intake prompts prophages to lyse their bacterial hosts^9^.

Gut virome alterations have been linked to several disease states like inflammatory bowel diseases^6,7^, malnutrition^10^, and type 2 diabetes^11^. But many such studies have not been able to identify specific viral lineages that are involved in such diseases, mainly due to the lack of viral marker genes^12,13^ and high phage diversity due to the rapid evolution^14^. Consequently, human gut phage studies are limited to relatively low taxonomic levels. While recent efforts uncovered viral families that are widespread in human populations, such as the crAss-like^15,16^ phages, these have not been successfully linked to disease states. In order to develop microbiome-targeted interventions to benefit human health, it is pivotal to study such higher-level phage taxonomies in the gut among relevant cohorts.

Here, we report on gut phageome alterations in metabolic syndrome (MetS)^17^ among 196 people. MetS is a collection of clinical manifestations that affects about a quarter of the world population, and is a major global health concern because it can progress into cardiometabolic diseases like type 2 diabetes, cardiovascular disease, and non-alcoholic fatty liver disease^18,19^. As gut bacteria are increasingly seen as contributing agents of MetS^20–22^, it stands to reason that the phages which infect these bacteria exhibit altered population compositions in MetS. For our analysis, we focused on dsDNA phages, which form a large majority of gut phages in particular and gut viruses in general^14,23^. We found MetS-connected decreases in phageome richness and diversity, which are correlated to bacterial population patterns. Such correlations extended to the relative abundance of phages and their particular bacterial hosts. We further identified eighteen viral clusters (VCs) that are significantly correlated with either MetS or healthy controls. We found that sequences contained in three of these VCs, one VC correlated with MetS and two VCs with controls, are related. These contain members of the *Candidatus Heliusviridae*, a previously unstudied lineage of temperate *Clostridiales* phages that is present in over 90% of the participants. Phylogenetic and taxonomic classification revealed at least six distinct *Ca. Heliusviridae* sub-groups, two of which are significantly more abundant in MetS. As the *Ca. Heliusviridae* include both phages which are associated with MetS and with healthy controls, this extremely widespread lineage is an interesting target for research into the human gut phageome.

## Results

### Metagenomic sequencing identifies high divergence in MetS phageomes

We performed bulk metagenome sequencing on fecal samples of 97 MetS and 99 healthy participants from the Healthy Life in an Urban Setting (HELIUS) cohort^24^, a large population study in Amsterdam, the Netherlands (**Supplementary Table 1**). This yielded an average of 42.1 ± 6.7 million reads per sample (median: 40.6 million reads). We assembled reads and selected 6,780,412 contigs longer than 1,500 bp or shorter but circular, among which we predicted phage sequences that we clustered at 95% nucleotide identity. This produced a database of 25,893 non-redundant phage contigs, which we grouped by shared protein content^25^ into 2,866 viral clusters (VCs). These comprised 14,433 contigs, while the remainder were singletons too distinct to confidently cluster with other phage contigs. Treating such singletons as VCs with one member gave a final dataset of 14,325 VCs.

Analysis of the read depth per VC across participants (**Supplementary .Table 2**) underscored the extremely high inter-individual diversity in gut phageomes, as 8,799 VCs (61.4% of total VCs) were specific to a single individual and 5,122 VCs (35.8% of total VCs) were found in two to twenty participants (*i.e.*, fewer than 10% of participants, **Supplementary Figure 1a**). Due to being so common, these two sets of VCs also comprised a mean of 92.6 ± 4.4% (median: 93.5%) of phage relative abundance (**Supplementary Figure 1b**). The remaining 241 VCs (1.7%) were present in over 10% of participants and represented 7.4 ± 4.4% (median: 6.6%) of phage relative abundance. Of these, 27 (0.2%) were found in over 30% of participants and may be part of the core human gut phageome^26^.

Next, we examined the relative abundance the of four VC sets (*i.e.*, individual-specific, present in ≤10, 10-30% and ≥30% of participants) in individual participants. For all four sets both the participant with the highest and lowest relative abundance of that VC set had MetS (**Supplementary Figure 1c**). This suggested greater β-diversity variation among MetS viromes than those in healthy controls, which we confirmed with a permutational analysis of multivariate dispersions (permdisp) on Bray-Curtis dissimilarities (*p* = 0.003, **Supplementary Figure 1d**). In conclusion, while we found high inter-individual diversity among phageomes in all participants, MetS phageomes exhibited greater β-diversity variation than controls.

### Gut phage and bacterial populations exhibit altered richness and diversity measures in MetS

To gain a deeper understanding of MetS phageome community dynamics, we first examined total read fractions that mapped to VCs. This was significantly lower in MetS compared to controls, implying that MetS participants either had lower phage loads or had higher absolute bacterial abundance than controls, or both (Wilcoxon signed rank test, *p* = 0.013, **Supplementary Figure 2a**). This pattern extended to prophage-carrying bacterial contigs, which likewise had lowered relative abundance among MetS participant than controls (Wilcoxon signed rank test, *p* = 1.4 × 10^−4^, **Supplementary Figure 2b**). This notably reflects a decrease in relative abundance of prophage-containing bacteria, not a decrease in that of temperate phages, as the relative abundances of VCs that encode the integrases used in prophage formation were unaltered (Wilcoxon signed rank test, *p* = 0.47, **Supplementary Figure 2c**). We hypothesize that prophage formation rates among MetS phageomes decrease and that phages possibly more commonly utilize the lytic phage life cycle. Furthermore, combined relative abundance of temperate VCs was a mean 34.8 ± 11.3% total phage relative abundance (median: 32.7%). Thus, while our bulk sequencing approach might be expected to bias in favor of prophages, the majority of phage relative abundance was composed of non-temperate phages.

For further analysis of phage communities, we examined phageome richness and diversity. We determined phage richness by measuring the number of VCs that were present (*i.e.*, had a relative abundance above 0) in each participant, using a horizontal coverage cutoff of 75%^27^. This showed that besides lower total phage relative abundance, MetS phageomes also had lower phage richness than controls, but equal evenness (Wilcoxon signed rank test, richness *p* = 8.7 × 10^−8^, Pielou evenness *p* = 0.79, **Figure 1a** and **b**). Nevertheless, due to the strong differences in species richness, phage α-diversity was significantly decreased among MetS participants (Shannon H’ *p* = 1.3 × 10^−3^, **Figure 1c**). This suggested that MetS gut phageomes are distinct from healthy communities. Indeed, MetS and control phageomes displayed significant separation when assessed by principal covariate analyses (PCoA) of β-diversity based on Bray-Curtis dissimilarities (permanova *p* = 0.001, **Figure 2d**).

**Figure 1:**
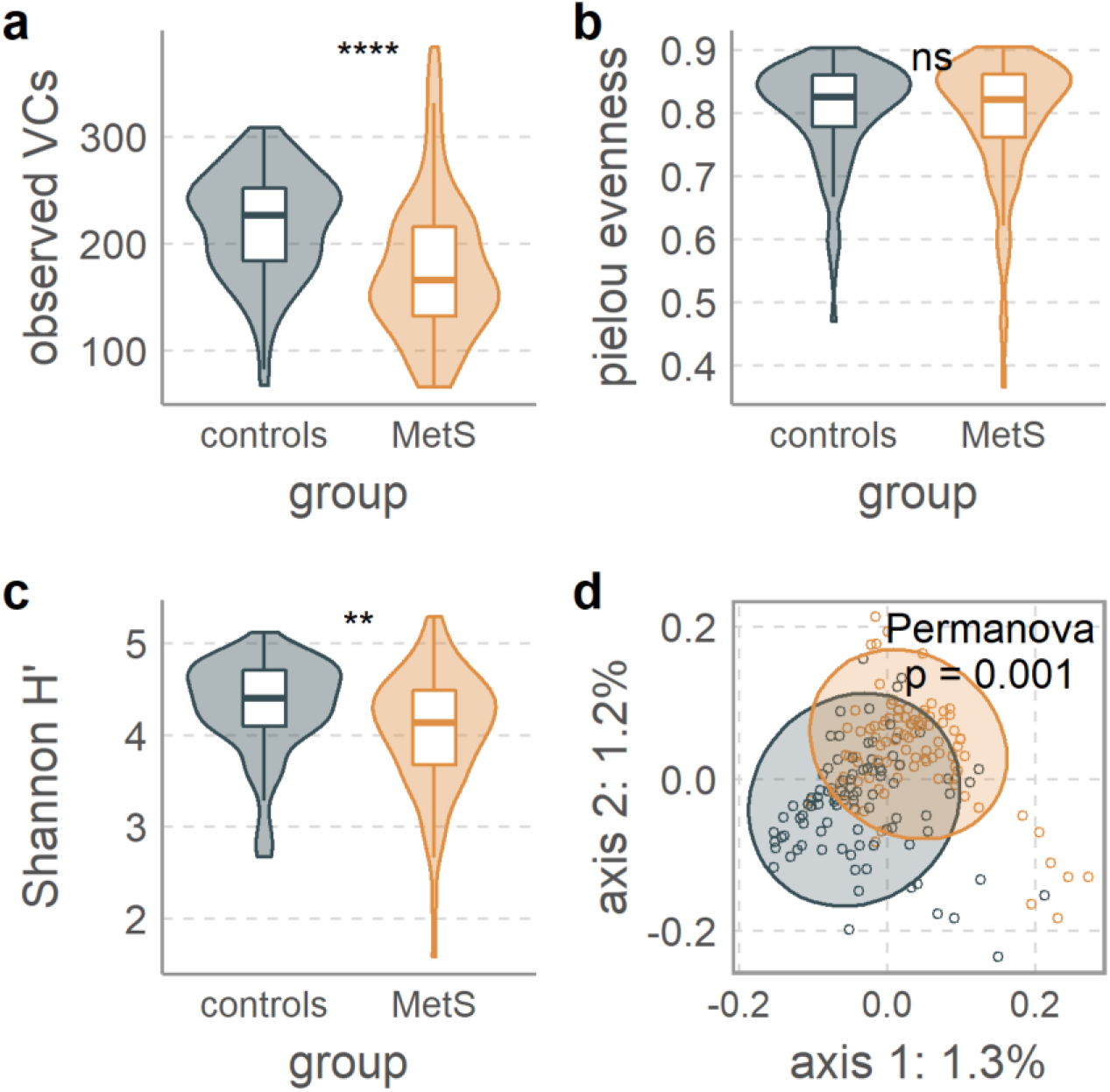
Gut phage populations are altered in MetS. **a** MetS-associated decreased phage species richness is evidenced by the number of unique VCs observed per sample. **b** No change in phage pielou evenness measurements. **c** significantly decreased phage α-diversity measured by Shannon diversity. **d** clear separation between phageomes of MetS and control participant as shown by β-diversity depicted in a principal coordinates analysis (PCoA) of Bray-Curtis dissimilarities. Permanova test was adjusted for smoking, age, sex, alcohol use, and metformin use. Statistical significance in A-C is according to the Wilcoxon signed rank test, where p-values are denoted as follows: ns not significant, * ≤ 0.05, ** ≤ 0.01, *** ≤ 0.001, **** ≤ 0.0001. Box plots show the median, 25^th^, and 75^th^ percentile, with upper and lower whiskers to the 25^th^ percentile minus and the 75^th^ percentile plus 1.5 times the interquartile range.

**Figure 2:**
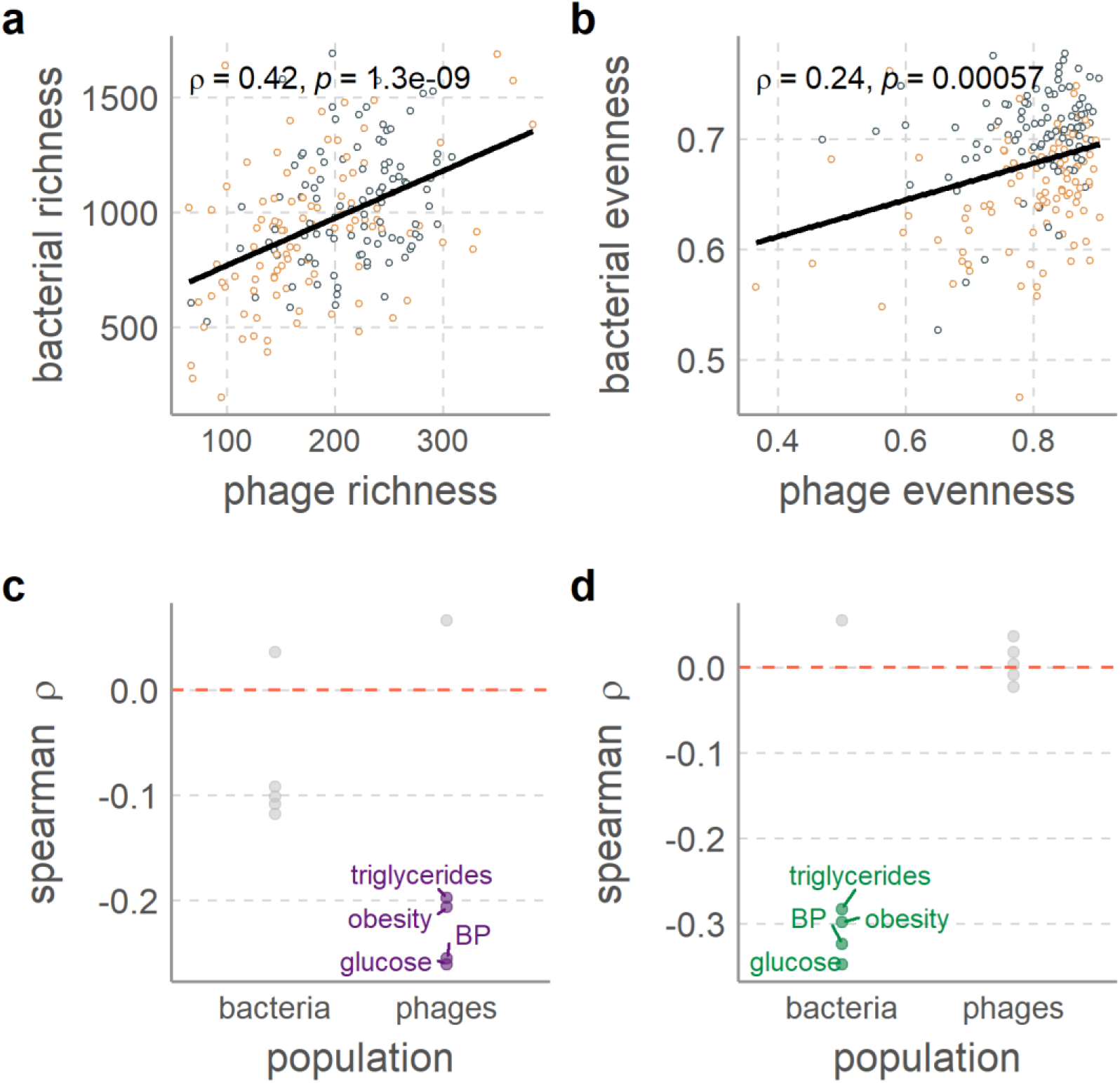
Correlations between phage and bacterial populations as well as between population measures and MetS clinical parameters. Strong correlations between **a** phage richness (observed VCs) and bacterial richness (Chao1 index), as well as between **b** phage and bacterial evenness (Pielou’s index), both with significant positive Spearman’s rank correlation coefficient. Both of these measures were correlated to MetS clinical parameters. Plotted are the Spearman’s rank correlation coefficients between the five MetS risk factors and **c** richness and **d** evenness. Points with p-values below 0.05 are colored in and labeled.

Because phages are obligate parasites of bacteria, we also studied bacterial 16s rRNA amplicon sequencing data. This showed that MetS phageomes mirror bacterial communities in species richness and α-diversity, but not evenness, which was significantly lowered in MetS bacterial populations (Wilcoxon signed rank test, Chao1 richness *p* = 9.1 × 10^−4^, Shannon *H’ p* = 1.5 × 10^−15^, Pielou evenness *p* =1.8 × 10^−14^, **Supplementary Figure 3a-c**). Additionally, bacterial communities separated in PCoA analysis in similar fashion to phageomes (permanova *p* = 0.001 for both bacteria and phages, **Supplementary Figure 3d**). Population-level phageome changes in MetS are thus directly related to a depletion of host bacteria populations, an assertion strengthened by significant direct correlations between phage and bacterial communities in richness (Spearman ρ = 0.42, *p* = 1.3 × 10^−9^, **Figure 2a**), evenness (Spearman ρ = 0.24, *p* = 5.7 × 10^−4^, **Figure 2b**).

As the bacterial and phage populations did not equally decrease in richness and evenness, they also did not equally correlate with MetS clinical parameters. Rather, phage richness was significantly negatively correlated with obesity, blood glucose levels, blood pressure, and triglyceride concentrations but bacterial richness was not (*p* < 0.05, **Figure 2c**). Bacterial evenness, meanwhile, did significantly negatively correlate with these clinical parameters while phage evenness did not (*p* < 0.05, **Figure 2d**). Increasingly severe MetS phenotypes thus result in stronger decreases in bacterial evenness than richness, while phage populations exhibit stronger decreases in richness than evenness. The decreasing bacterial evenness could be caused by depletion of certain bacterial species in MetS, which results in the viruses infecting these depleted bacteria to become undetectable, thereby decreasing richness more than evenness. Otherwise, the success of certain bacterial species could also decrease evenness. In the process this could obfuscate rare phage species, which could cause the decreased phage richness. Combined with the results showing MetS-associated reduction in total phage abundance but no increase in lysogeny (**Supplementary Figure 2**), our findings indicate that certain phages are either completely removed from the gut or become too rare to detect in MetS.

### Phages infecting select bacterial families are more abundant in MetS phageomes

We next studied individual bacterial lineages and the phages that infect them. To do this, we linked viral contigs to bacterial hosts by determining CRISPR protospacer alignments, taxonomies of prophage-containing bacterial sequences, and hosts of previously isolated phages co-clustered in VCs (see methods for details). We found 2,621 host predictions between 2,575 VCs (18% of all VCs) and eleven bacterial phyla, most commonly *Firmicutes* (2,067 VCs) and *Bacteroidetes* (234 VCs, **Supplementary Table 3**). We also identified 43 VCs with multi-phyla host range predictions, similar to previous works^28^.

To increase statistical accuracy, we selected 1,744 host predictions between 1,514 VCs (10.6% of all VCs) and the twelve most commonly occurring host families. We then performed an analysis of compositions of microbiomes with bias correction (ANCOM-BC)^29^, which showed higher relative abundances in controls for *Ruminococcaceae* VCs (*q* = 9.1 × 10^−3^), and in MetS for *Bacteroidaceae* VCs (*q* = 2.2 × 10^−4^), plus marginally for *Acidaminococcaceae* and *Tannerellaceae* VCs (*q* = 0.04, **Figure 3a**). ANCOM-BC on 16s rRNA gene sequencing data found that multiple *Ruminococcaeae* ASVs were significantly differentially abundant in controls (**Supplementary Figure 5**). Of *Ruminococcaceae* VCs with host predictions at the species level, those linked to *Faecalibacterium sp. CAG:74*, *Ruthenibacterium lactatiformans,* and *Subdoligranulum sp. APC924/74* had higher relative abundance in controls (Wilcoxon signed rank test with Benjamini and Hochberg adjustment, *q* ≤ 0.05, **Figure 3b**). These results are congruent with *Ruminococcaeae* being commonly linked to healthy gut microbiomes^3,30,31^.

**Figure 3:**
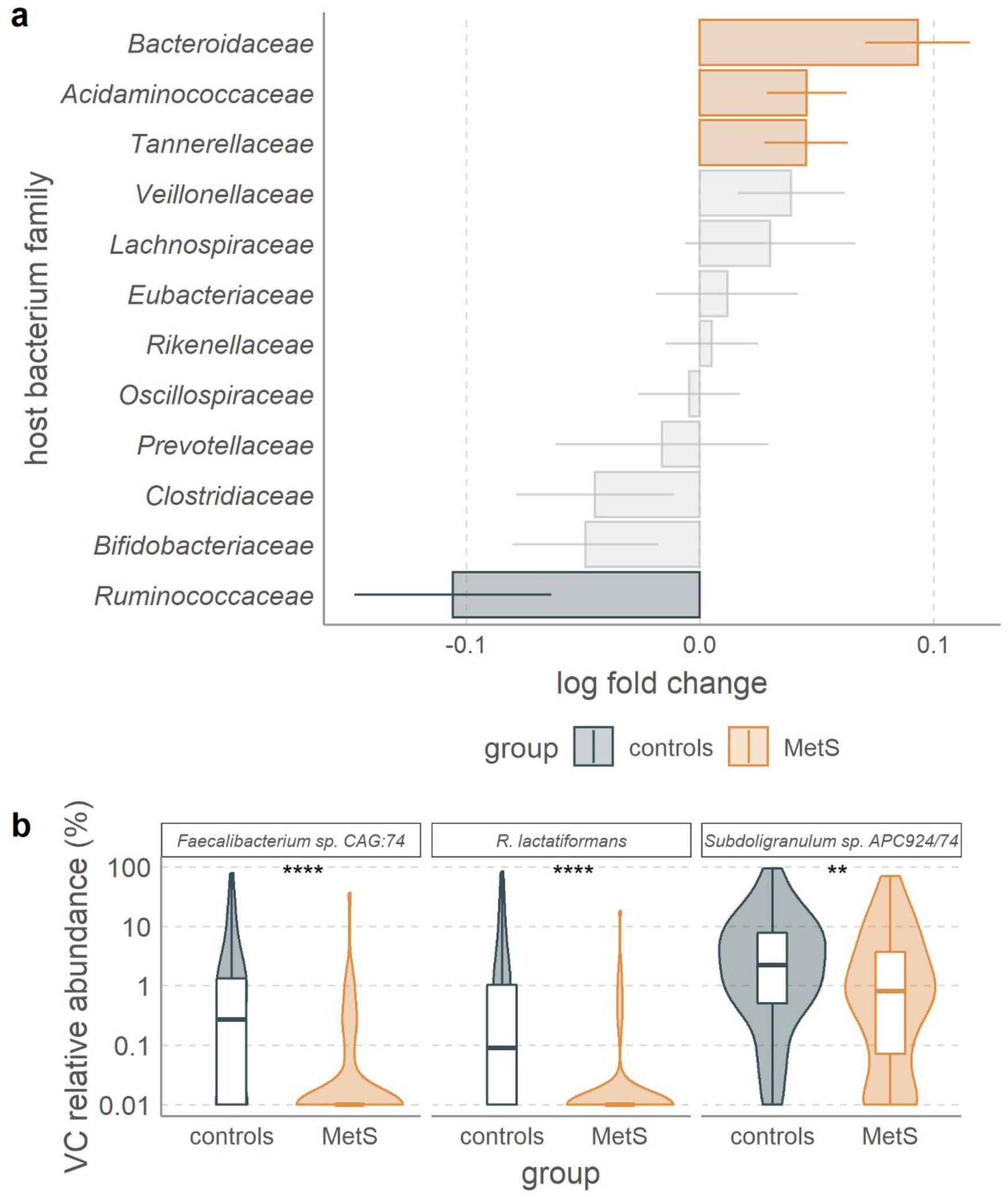
Phages infecting selected bacterial families are differentially abundant in MetS or healthy controls. **a** ANCOM-BC^29^ analysis of VCs that infect the twelve bacterial families to which the most VCs were linked shows significant association between *Bacteroidaceae* VCs and MetS, as well as between *Ruminococcaceae, Acidominacoccaceae, and Tannerellaceae* VCs and healthy controls. ANCOM-BC was adjusted for smoking, age, sex, alcohol use, and metformin use. **b** relative abundance comparisons between MetS and control participants of VCs infecting *Faecalibacterium sp.* CAG:74, *Ruthenibacterium lactatiformans*, *Subdoligranulum sp.* APC924/74. Stars denote significance according to the Wilcoxon signed rank test, with p-values adjusted with the Benjamini and Hochberg procedure (*q*). * ≤ 0.05, ** ≤ 0.01, *** ≤ 0.001, **** ≤ 0.0001. Box plots show the median, 25^th^, and 75^th^ percentile, with upper and lower whiskers to the 25^th^ percentile minus and the 75^th^ percentile plus 1.5 times the interquartile range. Error bars in **a** denote the standard error adjusted by the Benjamini-Hochberg procedure for multiple testing.

ANCOM-BC on 16s rRNA sequencing data identified *Bacteroidaceae* bacteria as significantly differentially abundant among MetS participants (*q* = 1.03 × 10^−13^). Since some widespread crAss-like gut phages infect *Bacteroidaceae* hosts^32,33^, we ascertained whether this phage family was more abundant among MetS participants. We did not find significant relations between crAss-like phage VC relative abundance and MetS (**Supplementary Figure 4a**), although VCs containing such phages were present more often among control (70/99) than MetS (57/97) participants (Fisher’s exact test, *p =* 0.1, **Supplementary Figure 4b**). Next to being absent more often, the participant with the highest relative abundance of crAss-like phage VCs belonged to the MetS group (17.2% of total phage relative abundance, **Supplementary Figure 4a**), which was indicative of greater variation in crAss-like phage relative abundance among MetS (mean 1.29±2.62%) than controls (mean 0.830±1.44%). MetS-associated alterations to crAss-like phage composition may thus occur at an individual level.

### *Bacteroidaceae* VCs are markers of the MetS phageome

The above results all indicate that MetS gut phageomes are distinct from those in healthy individuals. In light of this, we surveyed our cohort for individual VCs that were correlated with either MetS or healthy control phageomes. ANCOM-BC uncovered thirty-nine VCs that were more abundant in MetS participants, and eight more in controls (*q* ≤ 0.05, **Figure 4a**).

**Figure 4:**
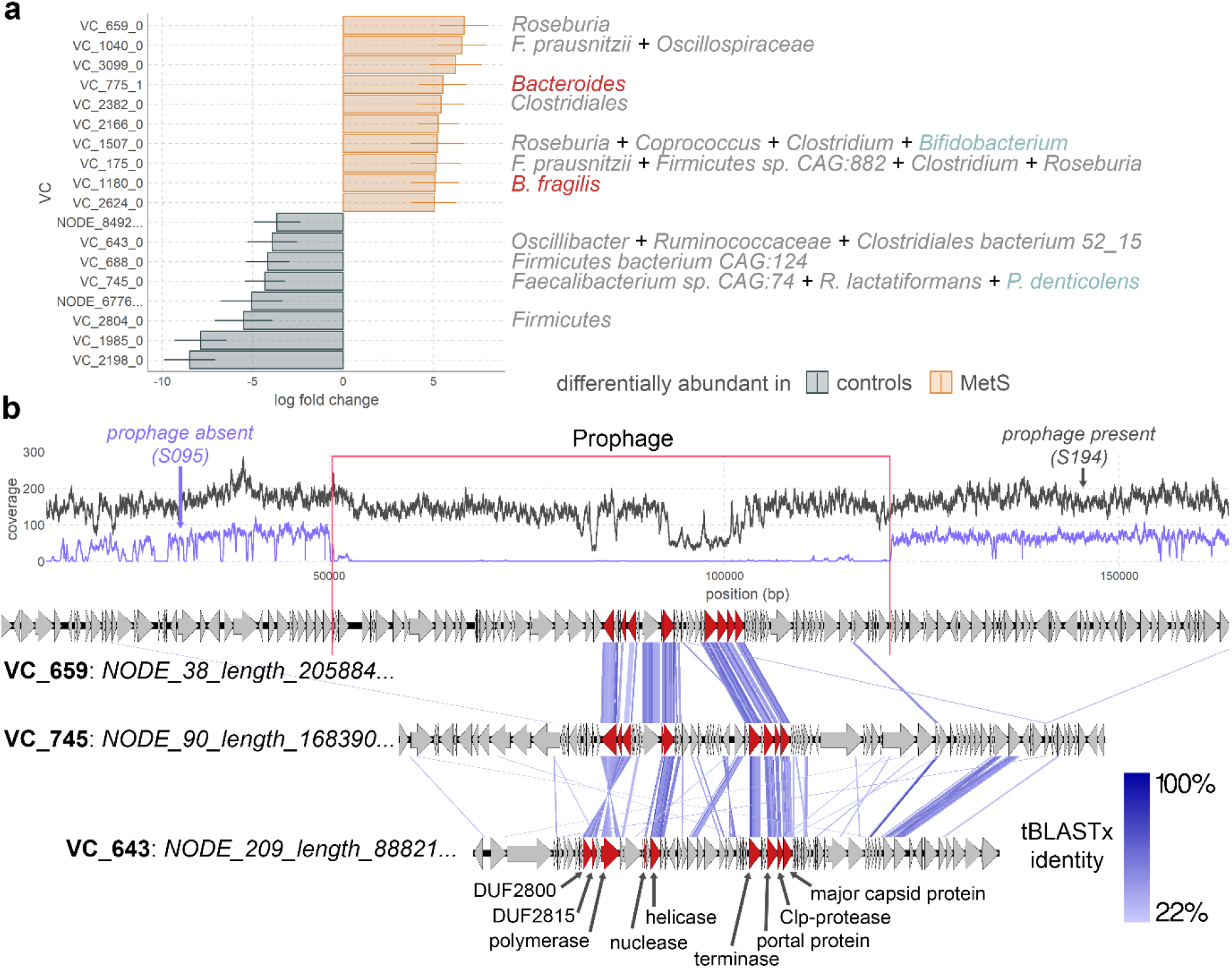
Among significantly differentially abundant VCs some are related. **a** VCs identified by ANCOM-BC analysis as significantly abundant (q ≤ 0.05 after implementing the Benjamini-Hochberg procedure for multiple testing). Error bars denote ANCOM-BC-supplied standard error. The analysis was adjusted for smoking, age, sex, alcohol use, and metformin use. Taxonomic names to the right of the plot denote host predictions, which are colored as follows: *Firmicutes;* grey, *Bacteroidetes;* red, *Actinobacteria*; green. The full taxonomies are listed in Supplementary Tables 2 and 4. For brevity, only the ten VCs most significantly associated with MetS (out of 38) are shown. See Supplementary table 7 for a full reckoning of significant VCs and the full names of the two singletons. **b** Whole genome analysis of three contigs that belong to VC_659_0, VC_745_0 and VC_643_0. The VC_659_0 contig is zoomed in on the prophage region, for the entire contig, see Figure 6. The read coverage depth of this contig in two samples is displayed at the top, on in which the prophage is present (S194) and one in which it is absent (S095). The nine genes shared by all *Candidatus Heliusviridae* are colored red, and annotated at the bottom.

In line with the above findings that *Bacteroidaceae* VCs are hallmarks of the MetS phageome, three MetS-associated VCs infected *Bacteroides* bacteria. The first (VC_1180_0) contained a non-prophage contig (*i.e.*, no detected bacterial contamination) of 34,170 bp with a checkV^34^ completion score of 100%. It further co-clustered with a contig that checkV identified as a complete prophage flanked by bacterial genes. Analysis with the contig annotation tool (CAT^35^) identified this contig as *Bacteroides fragilis*. Additionally, the most complete VC_1180_0 contig shared 6/69 (8.7%) ORFs with *Bacteroides uniformis Siphoviridae* phage Bacuni_F1^36^ (BLASTp bit-score ≥ 50). The second MetS-associated *Bacteroides* VC (VC_786_0) contained one contig with CRISPR spacer hits to *Bacteroides*. Its most complete contig had a CheckV completeness score of 98.94% and was classified by the contig annotation tool (CAT^35^) as *Phocaeicola vulgatus* (formerly *Bacteroides vulgatus*^37^). This near-complete contig furthermore shares 11/77 ORFs (14.3%) with *Riemerella Siphoviridae* phage RAP44 (BLASTp bit scores >50). This last finding was notable because the third and final MetS-associated *Bacteroides* VC (VC_775_1) also contained a near-complete genome (CheckV: 90.32% complete) that shared 16/81 ORFs (19.8%) with RAP44. Comparison of the most complete VC_786_0 and VC_775_1 contigs indicated that they share nine ORFs, revealing that they are part of an extended family of *Bacteroidetes Siphoviridae* phages of which members are hallmarks of MetS.

### A widespread phage family contains markers for healthy and MetS phageomes

Besides the above-mentioned *Bacteroidaceae* VCs, all other differentially abundant VCs with host links, two MetS- and four control-associated, infected *Firmicutes*, particularly in the *Clostridiales* order. The sole exception to this (VC_745_0) remarkably had CRISPR protospacer matches to *Firmicutes* bacteria *Faecalibacterium sp.* CAG: 74_58_120 and *Ruthenibacterium lactatiformans,* as well as to *Actinobacteria* bacterium *Parascardovia denticolens*. As this VC included genome fragments with simultaneous CRISPR protospacer hits to both phyla, VC_745_0 seemingly contains phages with an extraordinarily broad host range.

Besides this broad host range VC, our attention was drawn to the two MetS-associated *Clostridiales* VCs. Both were predicted to infect hosts that are usually associated with healthy gut microbiomes: *Roseburia*^3^ for VC_659_0, and *Oscillospiraceae*^38^ and *Faecalibacterium prausnitzii*^39^ for VC_1040_0. Further examination of their largest genomes revealed that MetS-associated VC_659_0 was remarkably similar to two VCs that were significantly associated with healthy controls: the above-mentioned broad host-range VC_745_0 and the less broad host-range *Oscillibacter/Ruminococcaceae* VC_643_0 (**Figure 4b**).

Intrigued by this apparent relatedness of VCs that included markers of MetS and healthy controls among our cohort, we sought to identify additional related sequences among our cohort. For this we first determined the exact length of VC_659_0 genomes by analyzing read coverage plots of a prophage flanked by bacterial genes (**Figure 4b**). By analyzing coverage of the contig in subjects where bacterial genes were highly abundant but viral genes were absent, we extracted a genome of 68,665 bp long. Homology searches of all 74 ORFs encoded by this prophage against all ORFs from all phage contigs in the cohort identified 249 contigs of over 30,000 bp that all shared nine genes (BLASTp bit score ≥ 50, **Figure 4b**). Additionally, we identified 51 *Siphoviridae* phage genomes in the National Center for Biotechnology Information (NCBI) nucleotide database that also shared these nine genes. With one exception, these were *Streptococcus* phages, the exception being *Erysipelothrix* phage phi1605.

The genes shared by all these phage genomes formed three categories. First are genes encoding structural functions: a major capsid protein, portal protein, CLP-like prohead maturation protease, and terminase. The second group are transcription-related genes encoding a DNA polymerase I, probable helicase, and nuclease. Finally, there are two genes that encode domains of unknown function, but which given their adjacency to the second group are likely transcription-related.

Earlier studies have used a cutoff of 10% gene similarity for phages that are in the same families, 20% for sub-families, and 40% for genera^40,41^. The nine shared genes form 10-25% of ORFs found on both the characterized phages and non-provirus contigs with checkV ‘high-quality’ designations. Thus, these phages form a family, which we dubbed the *Candidatus Heliusviridae*. Next, we further studied the *Ca. Heliusviridae* interrelatedness by calculating the pairwise percentages of shared protein clusters and hierarchically clustering the results (**Figure 5a**. This showed that the *Ca. Heliusviridae* form six groups, which we designated as *Ca. Heliusviridae* group alpha, beta, gamma, delta, epsilon, and zeta *Heliusvirinae*. A concatenated phylogenetic tree made from alignments of nine conserved *Ca. Heliusviridae* genes largely confirmed the hierarchical clustering (**Supplementary Figure 6**).

**Figure 5:**
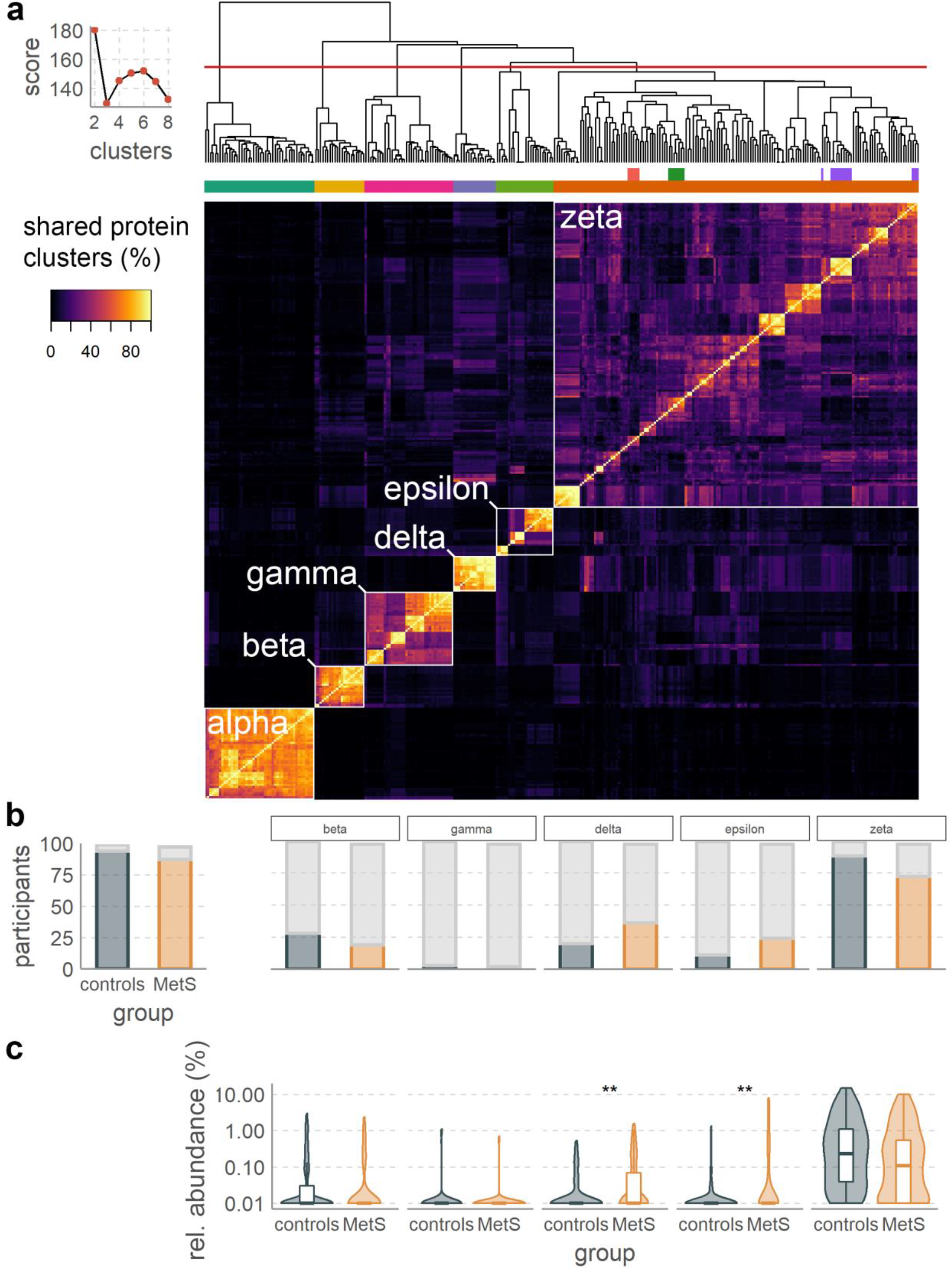
Three VCs that are hallmarks for either MetS or healthy control phageomes are part of the widespread *Candidatus Heliusviridae* family of gut phages. **a** heatmap and hierarchical clustering of pairwise shared protein cluster values for 249 contigs from the current study and 51 previously isolated phages. The line graph shows the optimal number of clusters as determined using the NbClust R package^109^, and the dendrogram is cut to form six clusters. These six clusters are labeled as alpha, beta, gamma, delta, epsilon, and zeta subfamilies. The top row of colors beneath the dendrogram denote the differentially abundant VCs, from left to right: VC_745_0 (red), VC_659_0 (green), and VC_643_0 (purple). The bottom colors are according to the candidate subfamilies. **b** the prevalence of the *Candidatus Heliusviridae* (left) and the separate candidate subfamilies (right). **c** the relative abundances of the candidate subfamilies (the whole family was not significantly more abundant in either group and is thus not depicted). q-values are denoted as follows * ≤ 0.05, ** ≤ 0.01, *** ≤ 0.001, **** ≤ 0.0001. Box plots show the median, 25^th^, and 75^th^ percentile, with upper and lower whiskers to the 25^th^ percentile minus and the 75^th^ percentile plus 1.5 times the interquartile range.

The *Ca. Heliusviridae* group alpha solely contained previously isolated *Streptococcus* phages, which both in the hierarchical clustering and the phylogenetic tree were distinct from the other genomes. Meanwhile, all three VCs that were significantly associated with either MetS or healthy controls where part of the *Ca. Heliusviridae* group zeta, by far the largest and most diverse group. Two of these, VC_659_0 and VC_745_0, formed distinct sub-clades in both hierarchical clustering and phylogenetic tree, while VC_643_0 conversely was spread out over multiple clades.

The *Ca. Heliusviridae* were present in 181 participants (92.3%), 94 controls and 87 MetS participants (**Figure 5b**). We also tested this finding in two cohorts in which the gut phageome was studied earlier, in the context of hypertension^42^ and type 2 diabetes^11^. To allow for incomplete assemblies, we searched for contigs in these two cohorts that contain the four conserved *Ca. Heliusviridae* structural genes. A phylogenetic tree containing concatenated alignments of the structural genes clearly showed that both validation cohorts contained sequences from all *Ca. Heliusviridae* groups (**Supplementary Figure 7**). Only a small minority of 47 sequences, largely from the hypertension cohort, formed a separate and distant clade of which the relation to the remainder of *Ca. Heliusviridae* is unclear. Among the two cohorts, *Ca. Heliusviridae* were present in 140/196 (71.4%, hypertension) and 112/145 (77.2%, T2D) participants. Finally, as this study and the two validation cohorts all utilized whole genome shotgun sequencing, the phages identified here might be inactive prophages. Thus, we used datasets of fecal virus-like particle (VLP) sequencing from ten people that were published earlier^43^. Cross-assembly of the ten VLP sequence datasets identified one contig of 43,244 bp (70.68% checkV completeness) and eight contig fragments that contained four or more conserved *Ca. Heliusviridae* genes. Thus, phages in this family are also found in VLP fractions, implying that they are inducible.

Of the *Heliusviridae* groups, the zeta was by far the most abundant, being present in 88 controls and 72 MetS participants. This meant healthy control phageomes were significantly more likely to contain *Heliusviridae* group zeta (Fisher exact test, *p =* 0.0096), though they were not significantly more abundant. Of the other candidate sub-families, the groups delta and epsilon were in significantly higher relative abundance (Wilcoxon signed rank test, *p* = 0.0043 and 0.0063, respectively) among MetS participants. The *Ca. Heliusviridae* group delta infects *Lachnospiraceae*, in particular *Butyrivibrio* sp. CAG:318 and *Lachnoclostridium* sp. An181. Meanwhile, the *Heliusviridae* group epsilon were distinct among the *Heliusviridae* in that they infect *Negativicutes* rather than *Clostridia*, specifically *Acidominacoccus* and several other genera in the *Veillonellaceae* (**Supplementary Table 3**). These results, combined with the fact that group zeta VC_659_0 is strongly correlated with MetS (**Figure 4a**), show that Ca. *Heliusviridae* are part of the core human gut phageome, where they may affect intricate relations with human health.

### MetS-associated group zeta *Ca. Heliusviridae* prophages encode possible metabolic genes

The *Ca. Heliusviridae* are generally linked to bacteria that are associated with healthy human gut microbiomes. Therefore, it is an apparent contradiction that MetS-associated *Ca. Heliusviridae* group zeta VC_659_0 infected *Roseburia*, a short chain fatty acids producer that is often abundant in healthy microbiomes^44^. Due to this contradiction, we explored this VC further. It contained two additional prophages, which where both incomplete (**Figure 6a**). Whole genome alignment showed that all three prophages shared their phage genes, and that the two incomplete ones also shared host-derived genes. This indicated that the incomplete prophages integrated into highly similar host bacteria which were distinct from *Roseburia*. To confirm this, we performed homology searches of the bacterial host ORFs found on these contigs against the NCBI nr database (BLASTp, bit-score ≥50). In both incomplete prophages, the majority of ORFs had *Blautia* as their top hit, which for a plurality of ORFs involved *Blautia wexlerae* (**Figure 6a**). VC_659_0 thus contains MetS-associated phages that integrate into at least two genera (*Roseburia* and *Blautia*) within the *Lachnospiraceae*.

**Figure 6:**
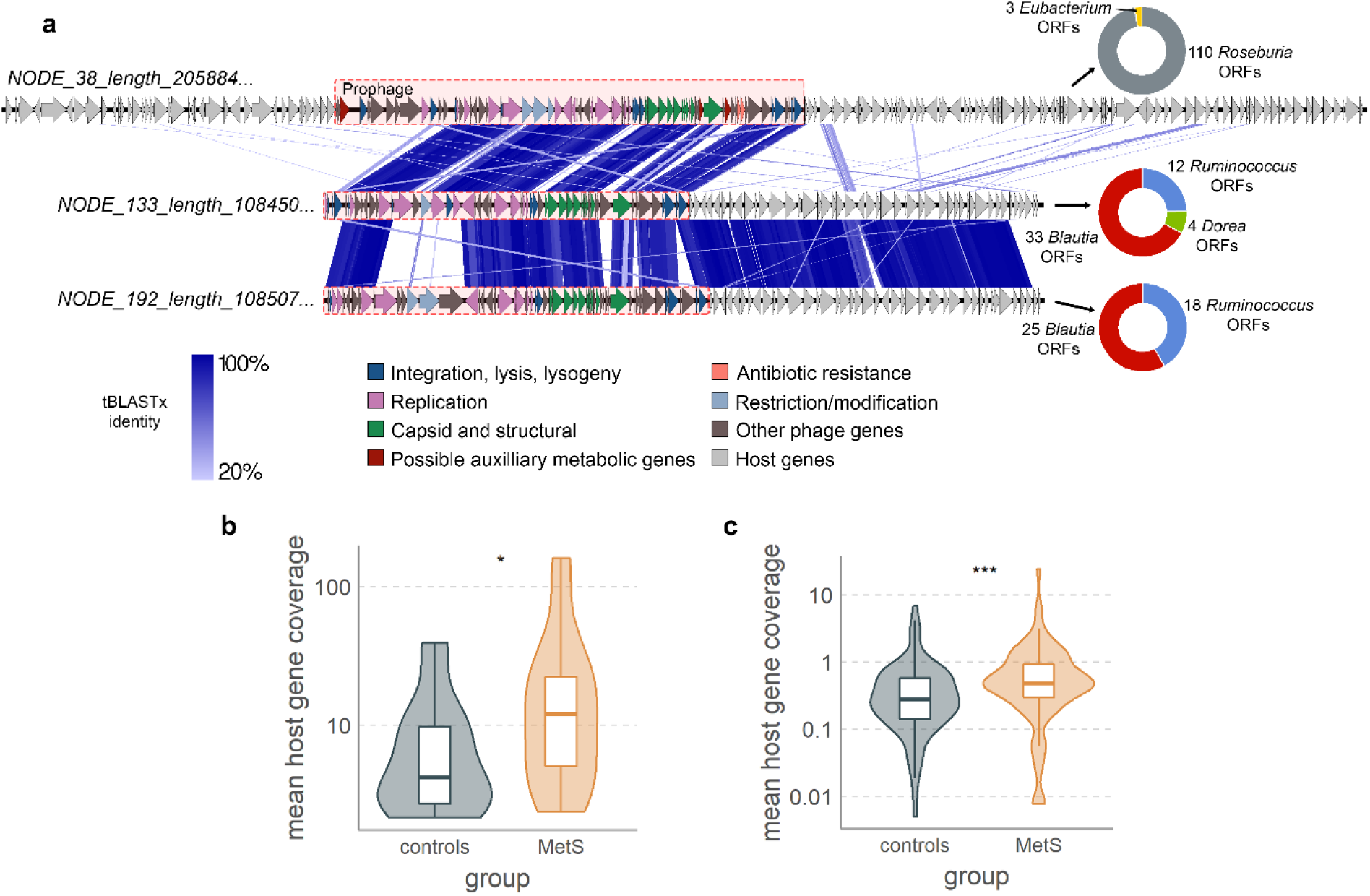
VC_659_0 infects *Roseburia* and *Blautia*, and carries possible auxiliary metabolic genes. **a** Whole genome alignment of three prophages contained within VC_659_0, with pie charts denoting the top BLASTp hit of all host genes on the contigs. **b** and **c** the mean coverage of host-derived regions in NODE_38 (**b**) and NODE_192 (**c**). Significance according to Wilcoxon signed rank test, p-values are denoted as follows * ≤ 0.05, ** ≤ 0.01, *** ≤ 0.001, **** ≤ 0.0001. Box plots show the median, 25^th^, and 75^th^ percentile, with upper and lower whiskers to the 25^th^ percentile minus and the 75^th^ percentile plus 1.5 times the interquartile range.

To examine if the hosts infected by VC_659_0 were more abundant in MetS participants, we determined mean coverage of bacterial genes found adjacent to the prophages. We thus assured that we analyzed the particular host strains infected by these phages, rather than unrelated strains in the same genera. This showed that both the *Blautia* and the *Roseburia* host genes were more abundant among MetS participants (Wilcoxon signed rank test, *Blautia p* = 5.1 × 10^−4^, *Roseburia p* = 0.042, **Figure 6b** and **c**). The specific *Lachnospiraceae* strains infected by VC_659_0 phages thus seem to thrive in MetS microbiomes. This could in part be due to functions conferred upon these bacteria by these prophages, as particularly the *Roseburia* prophage which carried several virulence and metabolism-related genes, including ones encoding a chloramphenicol acetyltransferase 3 (2.3.1.28), Glyoxalase/Bleomycin resistance protein (IPR004360), multi antimicrobial extrusion protein (IPR002528), 2-succinyl-6-hydroxy-2,4-cyclohexadiene-1-carboxylate synthase (4.2.99.20), and NADPH-dependent FMN reductase (PF03358). The latter two in particular are both associated with vitamin K (menaquinone) metabolism, which is part of (an)aerobic respiration in bacteria^45^. We speculate that this opens up the possibility that this *Roseburia* prophage aids its host bacterium, which in turn may contribute to MetS phenotypes.

## Discussion

This is the first study of the gut phages in the context of MetS, a widespread global health concern to which the gut bacteria targeted by phages are believed to be a main contributor^18^. We have shown that MetS is associated with decreases in gut phageome total relative abundance and richness, but no change in evenness. Due to their compositional nature, these phageome alterations could be bacterially driven, as phage total relative abundance decreases could be caused by bacterial counts increasing rather than phage counts decreasing. But since we measured decreased bacterial richness and evenness, MetS gut metagenomes would need to have larger numbers of bacterial cells that are distributed among fewer strains that are more unevenly divided than in healthy individuals. Conversely, total phage relative abundances could be lower in MetS due to lower viral loads, which would be in line with decreased phage richness and is in agreement with recently reported direct correlations between gut viral and bacterial populations in healthy individuals^46^. Future confirmation of this would necessitate counts of viable bacterial cells and VLP. In either case, we surmise that the main driver of these effects is diet, which affects bacterial^47–49^ as well as viral^50^ populations. It is also possible that phage populations as described here may further exacerbate bacterial diversity losses, as low phage abundance may decrease their positive effects on bacterial diversity^51,52^.

One aspect in which our study separated itself from some other gut virome studies was its usage of whole genome rather than VLP sequencing. We believe our approach has its advantages, such as a lack of pre-sequencing amplification and the biases this introduces, and a greater emphasis on the prophage community significant to functioning of both bacterial and phage communities in the gut^8,9,12,53^. It must nevertheless be noted that our sequencing approach may underestimate the virulent proportion of the human gut phageome, likewise to VLP sequencing approaches probably underestimating gut prophage communities. Indeed, analysis of both VLP and bulk communities is likely needed to gain a full reckoning of human gut phageomes^14^. This approach could furthermore distinguish between inducible and defective prophages^54^, which we were unable to do.

Like other studies, we found highly individual-specific^26,43^ human gut phageomes in which singular widespread phage VCs are rare^14,15^. Despite the universally high inter-individual phageome diversity, we found larger within-group variation among MetS phageomes than healthy controls. This is consistent with the Anna Karenina principle (AKP), which holds that “all healthy microbiomes are similar; each dysbiotic microbiome is dysbiotic in its own way”^55^. Such AKP-dynamics mirror previous findings of obesity-related alterations in the gut bacterial populations^56^. Particularly, we hypothesize that stressors inherent to MetS perturb gut bacterial populations in a stochastic fashion, the effects of which reverberate to the phage populations and result in their increased within-group variation.

We further found strong negative correlations between the risk factors that constitute MetS and phage richness but not evenness. This likely results from the whole-genome sequencing approach that we took, which better captures intracellular phages (*e.g.*, actively replicating or integrated prophages) than extracellular phages^14^. The phage VC richness that we report here thus represents phages that are actively engaging with their hosts or are highly abundant extracellularly. As phages that target depleted bacteria are more likely to be low in abundance and extracellular, our approach does not capture them. Thus, the apparent species richness drops because low abundant extracellular phages are below the detection limit of our sequencing approach. This removal of rare phages in turn prohibits significant drops in species evenness in MetS. It could also be that bacteria depleted in MetS reside in phage-inaccessible locales within the gut^57^, which perhaps results in removal of the corresponding phages from the gut to below detectable levels. This would explain the stronger correlation between bacterial evenness than richness to MetS risk factors.

As most (gut) phages remain unstudied^14,58^, it is often difficult to link phages to host bacteria^59^. Here, we linked roughly one tenth of all VCs to a bacterial host. The remaining majority of VCs likely represent phages that infect bacterial lineages lacking CRISPR systems^60^, are exclusively lytic, or that integrate into hosts which we could not taxonomically classify. Whichever is the case, our study underscores the great need for methods that link phages to hosts with high accuracy^61,62^. From the phage-host linkages that we obtained, we found that VCs containing phages infecting specific bacterial families tend to be either depleted (*Bifidobacteriaceae, Ruminococcaeae*, and *Oscillospiraceae)* or enriched (*Bacteroidaceae*) in tandem to their hosts. For the *Bifidobacteriaceae* and *Oscillospiracaeae*, this is in line with established studies that show depletion of these families in MetS^3^ and MetS-associated disease states^30,38^.

For *Ruminococcaceae* bacteria, associations with MetS and MetS-related diseases are less clear, with reports of both positive^3,31^ or negative^39,63^ associations. The specific *Ruminococcaceae* of which we link the phage-containing VCs to healthy controls, most notably *Faecalibacterium prausnitzii* and *Ruthenibacterium lactatiformans*, are both considered hallmarks of healthy human gut microbiomes^3,30,39^. Interestingly, we also succeeded in linking specific viral clusters that infect *Faecalibacterium* to both healthy controls (VC_745_0) and MetS participant groups (VC_1040_0). This contradiction may indicate that infections of *Faecalibacteria* could result in differing outcomes for the bacterium depending on the phage species. As both VCs contain sequences with integrase-like genes, they contain temperate phages. It could be that VC_745_0 prophages augment *Faecalibacterium* growth, as prophages are known to do^64^. Meanwhile VC_1040_0 prophages could be detrimental to the same bacteria, for example by becoming lytic in the presence of MetS-associated dietary components, likewise to *Lactobacillus reuteri* prophages lysing their hosts in the presence of dietary fructose^9^. Such behavior can lead to the collapse of the bacterial population^44^, and may thereby be a contributing factor to depletion of *Faecalibacterium* in MetS.

As mentioned, the *Bacteroidaceae* were the only bacterial family that are infected by phages of which the VCs were significantly more abundant among MetS participants. Concordantly, we found several individual *Bacteroides* VCs that were MetS-associated. The *Bacteroides* are often positively associated with high-fat and high-protein diets^65,66^. Simultaneously, however, reports disagree on individual *Bacteroides* species and their associations with MetS-related diseases like obesity, type 2 diabetes, and non-alcoholic fatty liver disease^30^. Such conflicting reports likely reflect the large diversity in metabolic effects at strain level among these bacteria^67^. Based on our results, we drew two conclusions. First, that *Bacteroidaceae*-linked VCs mirror their hosts in MetS-associated relative abundance increase, and second that *Bacteroidaceae*-linked VCs are of significant interest to studies of the MetS microbiome. The latter conclusion is strengthened by findings that *Bacteroides* prophages can alter bacterial metabolism in the gut^8^.

While *Bacteroidaceae* VCs at large were thus seemingly associated with MetS phenotypes, we uncovered larger variation of crAss-like phage-containing VC abundance, which suggest at individual-specific alterations to this gut phage family among MetS phageomes. This widespread and often abundant human gut phage family infects *Bacteroidetes*, including members of the *Bacteroidaceae*^68,69^. As these phages are commonly linked to healthy gut microbiomes^41,69,70^, it is conceivable that they would be negatively correlated with MetS phageomes. That this is not the case among the entire cohort is likely due to the great variety within this family^69^, and perhaps additionally to the hypothesized aptitude of crAss-like phages for host switching through genomic recombination^69^.

Finally, our study revealed the *Candidatus Heliusviridae*, a highly widespread family of gut phages that largely infect *Clostridiales* hosts. This prospective family is also expansive, and includes at least six distinct candidate subfamilies. Our uncovering of this novel human gut phage family underscores the usefulness of database-independent *de novo* sequence analyses^25,27,71^, as well as the need for a wider view on viral taxonomy than has presently been exhibited in the field of gut viromics.

The *Ca. Heliusviridae* are of particular interest to studies of MetS and related illnesses because its member phages include some associated with MetS and others with healthy controls. Most striking is the fact that most of the bacteria infected by MetS-associated *Ca. Heliusviridae*, are generally producers of short chain fatty acids (SCFA) such as butyrate and commonly depleted in MetS^30^. Such SCFA-producing bacteria are commonly positively associated with healthy microbiomes, as SCFAs that result from microbial digestion of dietary fibers have a role in the regulation of satiation^72,73^. The exception to this are the *Veillonellaceae* that are infected by the *Heliusviridae* group epsilon, which are found at elevated abundance in non-alcoholic fatty liver disease^30^. While higher abundance of some of the other butyrate-producers infected by *Ca. Heliusviridae* is associated with metformin use^74^, this is used to treat type 2 diabetes rather than MetS.

Particularly interesting are the *Roseburia/Blautia* phages in VC_659_0, which was the most strongly correlated with MetS out of all VCs. The positive correlation between the relative abundance of these phages and that of their hosts indicates that they have a stable relation with their hosts in the MetS microbiome. This is to be expected, as large-scale prophage induction is generally associated with sudden alterations to the microbiome, such as the addition of a specific food supplement that acts as an inducer of prophages^9^. Such sudden alterations in phage behavior are unlikely to be captured in large cohorts with single measurements. In fact, as phages are strongly dependent on their host, one might expect the abundance of many gut phages to be positively correlated to that of their particular hosts under the relatively temporally stable conditions of MetS. The strong correlation of VC_659_0 to MetS phenotypes, coupled to the commonly found correlation to healthy microbiomes of VC_659_0 host bacteria, and the presence of potential auxiliary metabolic genes in VC_659_0 phage sequences combined introduce the possibility that prophage formation of these *Ca. Heliusviridae* phages alters the metabolic behavior of their host bacteria, as is known to happen in marine environments^75,76^. This could make these bacteria detrimental to health. Proving this hypothesis necessitates future isolation of VC_659_0 phages.

Despite efforts to catalog the human gut phageome^14,28^, taxonomically higher structures are still largely absent. This study shows the worth of analyzing phages at higher taxonomic levels than genomes or VCs, similarly to what has been shown in recent years regarding the crAss-like phage family^15,16^. Unlike the crAss-like phage family, however, the *Ca. Heliusviridae* seem to be strongly correlated with human health. We hope that further research will provide a deeper understanding of the effect that these phages have on their bacterial hosts and the role that this plays in MetS, as well as a refinement of their taxonomy.

## Acknowledgements

PAdJ and TPMS were supported by a Senior Fellowship of the Dutch Diabetes Research Foundation (2019.82.004) to HH. KW was supported by a Novo Nordisk Foundation CAMIT grant 2018 to MN. BED was supported by the Netherlands Organization for Scientific Research (NWO) Vidi grant 864.14.004 and European Research Council (ERC) Consolidator grant 865694: DiversiPHI. The funders had no role in the study design, the collection, analysis, and interpretation of data, the writing of the report, and the decision to submit the article for publication.

The HELIUS study is conducted by the Amsterdam UMC, location AMC with the Public Health Service of Amsterdam. Both provided core financial support for HELIUS. The HELIUS study is also funded by research grants of the Dutch Heart Foundation (Hartstichting; 2010T084), the Netherlands Organization for Health Research and Development (ZonMw; 200500003), the European Integration Fund (EIF; 2013EIF013), and the European Union (Seventh Framework Programme, FP-7; 278901). We gratefully acknowledge the AMC Biobank for their support in biobank management and high-quality storage of collected samples. We are most grateful to the participants of the HELIUS study and the management team, research nurses, interviewers, research assistants and other staff who have taken part in gathering the data of this study.

## Author contributions

PAdJ and KW conducted data analysis; TPMS, BJvdB, AHZ, FLN, BED, and MN assisted with experimental design and data interpretation; PAdJ and HH designed the study and wrote the manuscript. All authors read and provided input on the manuscript.

## Declaration of Interests

MN owns stock in, consults for, and has intellectual property rights in Caelus Health. He consults for Kaleido. None of these are directly relevant to the current paper.

## Methods

### Sequencing and contig assembly

The Healthy Life in an Urban Setting (HELIUS) cohort includes some 25,000 ethnically diverse participants from Amsterdam, the Netherlands. The cohort details were published previously^77^. The HELIUS cohort conformed to all relevant ethical considerations. It complied with the Declaration of Helsinki (6^th^, 7^th^ revisions), and was approved by the Amsterdam University Medical Centers Medical Ethics Committee. For details on stool sample collection from among the participants, their storage, and DNA extraction, see Deschasaux, *et al*^24^. In summary, participants were asked to deliver stool samples to the research location within 6 hours after collection with pre-provided kit consisting of a stool collection tube and safety bag. If not possible, they were instructed to store their sample in a freezer overnight. Samples were stored at the study visit location at −20°C until daily transportation to a central −80°C freezer.

Total genomic DNA was extracted using a repeated bead beating method described previously^24,78^. Libraries for shotgun metagenomic sequencing were prepared using a PCR-free method at Novogene (Nanjing, China) on a HiSeq instrument (Illumina Inc. San Diego, CA, USA) with 150 bp paired-end reads and 6 Gb data/sample. All bioinformatics software was run using standard settings, unless otherwise stated. All sequencing reads are available at the European Nucleotide Archive under project PRJEB42542. Samples have accession numbers ERS5585222-ERS5585321, all phage contigs are under accession number ERZ1762427.

Following previously set definitions^79^, participants were classified in the MetS group if three of the following five health issues occurred: abdominal obesity measured by waist circumference, insulin resistance measured by elevated fasting blood glucose, hypertriglyceridemia, low serum high-density lipoprotein (HDL), and high blood pressure^79^. All participants of the HELIUS cohort reside in Amsterdam, the Netherlands. Participants were roughly evenly divided by ethnicity, with European Dutch comprising 49 controls and 49 MetS participants, and African Surinamese 50 controls and 49 MetS participants. The MetS group contained 55 women and had a median age of 58 (mean 56.8±8.09), and the controls 71 and had a median age of 50 (mean 49.1±12). Of the 196 participants, 26 used metformin, of whom 2 were controls who did not concur to the MetS criteria. Analysis of sequencing output started with assembly of the sequencing reads per sample (*i.e.,* 196 individual assemblies) into contigs using the metaSPAdes v3.14.1 software^80^. For each sample, we selected contigs of more than 5,000 bp for further analysis. In addition, among contigs between 1,500 and 5,000 bp we identified circular contigs by checking for identical terminal ends using a custom R script that employed the Biostrings R package v3.12^81^. All 6,780,412 circular contigs and contigs over 5,000 bp were then pooled before phage sequence prediction.

### Phage and bacterial sequence selection

We predicted phage sequences as described previously^82^. In short, we first analyzed contigs using VirSorter v1.0.6^83^ and selected those in category 1, 2, 4, and 5. In parallel, contigs were analyzed using VirFinder v1.1, after which we selected those with a score above 0.9 and a p-value below 0.05. We additionally classified contigs as phage if (I) they were both in VirSorter categories 3 or 6 and had VirFinder scores above 0.7 with p-values below 0.05, and (II) annotation with the contig annotation tool (CAT) v5.1.2^35^ was as “Viruses” or “unclassified” at the superkingdom level. After removing those with CAT classifications as Eukaryotic viruses, this resulted in a database of 45,568 phage contigs. Bacterial sequences were predicted by selecting all contigs that CAT annotated in the “Bacteria” at the superkingdom level, and removing contigs that were also found in the phage dataset. An exception was made for prophage contigs in VirSorter category 4, 5, and 6, which were left among the bacterial dataset (see “Phage-host linkage prediction”). This resulted in a total of 1,579,361 bacterial contigs. The 1,624,929 bacterial and phage datasets were then concatenated and deduplicated using dedupe from BBTools v38.84 with a minimal identity cutoff of 90% (option minidentity=90). This identified 759,403 duplicates and resulted in 829,633 non-redundant bacterial sequences and 25,893 non-redundant phage sequences. While the bacterial sequences were used for host prediction (see “Phage-host linkage prediction”), we subsequently predicted open reading frames (ORFs) in phage contigs using Prodigal v2.6.2^84^ (option -p meta). These ORFs were then used to group phage sequences in viral clusters (VCs) using vContact2 v0.9.18^25^. This resulted in 2,866 VCs comprising 14,433 phage contigs and 11,460 singletons and outliers, which we treated as VCs with one member. This resulted in 14,325 VCs. For a full accounting of phage contigs, see **Supplementary Table 2 and 4.**

### Read mapping and community composition

For bacterial community composition, we used sequencing data targeting the V4 region of the 16s rRNA gene that had been performed previously^24,85^. Details on ASV construction from these samples was described previously in Verhaar, et al^85^. As part of this previous analysis, samples with fewer than 5000 read counts had been removed, and samples had been rarified to 14932 counts per sample.

To determine phage community composition, we mapped reads from each sample to the non-redundant contig dataset using bowtie2 v2.4.0^86^. As previously recommended^27^, we removed spurious read mappings at less than 90% identity using coverM filter v0.5.0 (unpublished; https://github.com/wwood/CoverM, option-min-read-percent-identity 90). The number of reads per contig was calculated using samtools idxstats v1.10^87^. As was also recommended^27^, contig coverage was calculated with bedtools genomecov v2.29.2^88^, and read counts to contigs with a coverage of less than 75% were set to zero. Read counts for each sample were finally summed per VC. All contigs were analyzed for completion with CheckV v 0.7.0-1^34^.

### Ecological measures

In all boxplots, we tested statistical significance using the Wilcoxon rank sum test as it is implemented in the ggpubr v0.4.0 R package (available from: https://cran.r-project.org/web/packages/ggpubr/index.html). Unless stated otherwise, all plots were made using either ggpubr or the ggplot2 v3.3.2 R package (available from: https://cran.r-project.org/web/packages/ggplot2/index.html). Alpha diversity measures (observed VCs and Shannon H’ for phages and Chao1 and Shannon H’ for bacteria) were calculated using read count tables with the plot_richness function in the phyloseq R package v1.33.0^89^. For β-diversity, we converted read counts to relative abundances using the transform function from the microbiome v1.11.2 R package. We then used the phyloseq package to calculate pairwise Bray-Curtis dissimilarities and construct a principal coordinates analysis (PCoA). Statistical significance of separation in the PCoA analysis was determined with a permutational multivariate analysis of variance (permanova) using the adonis function from the vegan R package^90^. For this analysis, we adjusted for smoking, sex, age, alcohol use, and metformin use. Direct correlation coefficients between richness and diversity were calculated using the stat_cor function in the ggpubr R package.

### Phage-host linkage prediction

We predicted VC-bacterium links in three ways: (i) CRISPR protospacers, (ii) prophage similarity, and (iii) characterized phage similarity.

We predicted CRISPR arrays among the bacterial contigs using CRISPRdetect v2.4^91^ (option array_quality_score_cutoff 3) and used these to match bacterial contigs and phage contigs. In addition, we used a dataset of 1,473,418 CRISPR spacers that had previously been predicted^62,92^ in genomes contained in the Pathosystems Resource Integration Center (PATRIC)^93^ database to match to phage contigs with spacePharer v2-fc5e668^94^ using standard settings and cutoffs. This process resulted in 3,727 spacer hits, of which 2,244 hits were either to PATRIC genomes or to bacterial contigs from this study with definite CAT classifications at the phylum level (**Supplementary Table 3**).

To identify predicted phage contigs with high sequence similarity to prophages, we analyzed which viral clusters contained on of the 7,691 bacterial contigs with VirSorter prophage predictions in category 4 or 5. CAT was subsequently used to determine the taxonomy of bacterial contigs with prophage regions. In total, we linked 1,102 VCs to prophages with this approach.

Finally, VCs were linked to bacterial hosts by vContact2 clustering with characterized phages from the viral RefSeq V85 database^95^ with a known host. To achieve this, we selected all VCs from the vContact2 output that contained both characterized genomes and phage contigs. If all characterized phages infected hosts within the same bacterial family, we took that to mean that the whole VC infects hosts from that family. This approach linked 44 VCs to hosts.

### Differential abundance analysis

To determine which bacteria and VCs were differentially abundant between MetS and control subjects, we employed the analysis of composition of microbiomes with bias correction (ANCOM-BC)^29^. This novel method, unlike other similar methods like DeSeq2, takes into account the compositional nature of metagenomics sequencing data^96^. To implement this method, we applied the ANCOM-BC v1.0.2 R package to raw read count tables, as ANCOM-BC employs internal corrections for library size and sampling biases^29^. Significance cutoff was set at an adjusted p-value of 0.05, p-values were adjusted using the Benjamini-Hochberg method, and all entities (bacteria taxa/VCs) that were present in more than 10% of the samples were included (options p_adj_method = “BH", zero_cut = 0.9, lib_cut = 0, struc_zero = T, neg_lb = F, tol = 1e-5, max_iter = 100, alpha = 0.05). For this analysis, we adjusted for smoking, sex, age, alcohol use, and metformin use.

### crAss-like phages

To identify crAss-like phages, we employed a methodology described earlier^41^. Shortly, a BLAST database was made containing all ORFs from all phage contigs (predicted before viral clustering, see “Viral and bacterial sequence selection”) using BLAST v2.9.0+^97^. We then performed two BLASTp searches in this database, one with the terminase (YP_009052554.1) and one with the polymerase (YP_009052497.1) of crAssphage (NC_024711.1), with a bitscore cutoff of 50. All phage contigs that had (i) a hit against both crAssphage terminase and polymerase and a query alignment of ≥350 bp, and (ii) a contig length of ≥70 kbp were considered crAss-like phages. This resulted in 146 crAss-like phage contigs, which were contained in 29 VCs.

### Candidatus *Heliusviridae* analysis

To detect pairwise similarity, whole genome analyses were constructed with Easyfig v2.2.5^98^. The prophage borders in NODE_38_length_205884_cov_102.806990 were determined by determining the read depth along the entire contig from the bam files with read mapping data (“Read mapping and community composition”) using bedtools genomecov v2.29.2^88^ with option -bg. Resultant output was parsed and plotted in R. Other related phages among the cohort were detected by performing a BLASTp search with all phage ORFs of NODE_38_length_205884_cov_102.806990 against all phage ORFs of the cohort with Diamond v2.0.4. This identified nine genes that were present in 249 contigs. The ORFs on these contigs were annotated using PROKKA v1.14.6^99^ and InterProScan v5.48-83.0^100^. To identify isolated phages that share these nine contigs, we performed a BLASTp against the NCBI nr-database using the NCBI webserver^101^ on February 26 2021 and collected all genomes with hits against all nine genes (bit score ≥50).

The phages sharing all nine genes were clustered by analyzing them with vContact2 v0.9.18^25^, extracting the protein clustering data and calculating the number of shared clusters between each pair of contigs. Contigs were clustered in R based on Euclidean distances with the average agglomeration method.

To build a taxonomic tree, the nine genes were separately aligned using Clustal Omega v1.2.4^102^, positions with more than 90% gaps were removed with trimAl v1.4.rev15^103^ and alignments were concatenated. From the concatenated alignment, an unrooted phylogenetic tree was built using IQ-Tree v2.0.3^104^ using model finder^105^ and performing 1000 iterations of both SH-like approximate likelihood ratio test and the ultrafast bootstrap approximation (UFBoot)^106^. In addition, ten iterations of the tree were separately constructed, as has been recommended^107^ (Zhou et al., 2018) (IQ-Tree options -bb 1000, -alrt 1000, and --runs 10).

### Validation of *Ca. Heliusviridae* in other cohorts

We used three additional studies to analyze prevalence of the *Ca. Heliusviridae*; one composing of 145 participants used to study the gut virome in type 2 diabetes^11^, a second containing 196 participants and used to study the gut virome in hypertension^42^, and a final one containing ten healthy participants studied by VLP sequencing^43^. Reads belonging to all studies were downloaded from the NCBI sequencing read archive (SRA) and assembled as described above. The ten-patient VLP cohort was cross-assembled, while the other two cohorts were assembled separately. After assembly, ORFs were predicted using Prodigal v2.6.2^84^. *Ca. Heliusviridae* members were identified by blastp using Diamond v2.0.4^108^ against ORFs from each study, in which the terminase, portal protein, Clp-protease, and major capsid protein of NODE_38_length_205884_cov_102.806990 were used as queries. This was done instead of all nine signature *Ca. Heliusviridae* genes to better allow for incomplete assemblies. Contigs containing all four genes were selected, and a concatenated alignment was made of the four head genes found in the T2D and hypertension cohorts, plus all *Ca. Heliusviridae* in the tree depicted in Supplementary Figure 5. These were then used to build a phylogenetic tree. The concatenated alignment and phylogenetic tree were constructed as described above under “Candidatus Heliusviridae analysis”.

## Supplementary Figures

**Supplementary Figure 1:**
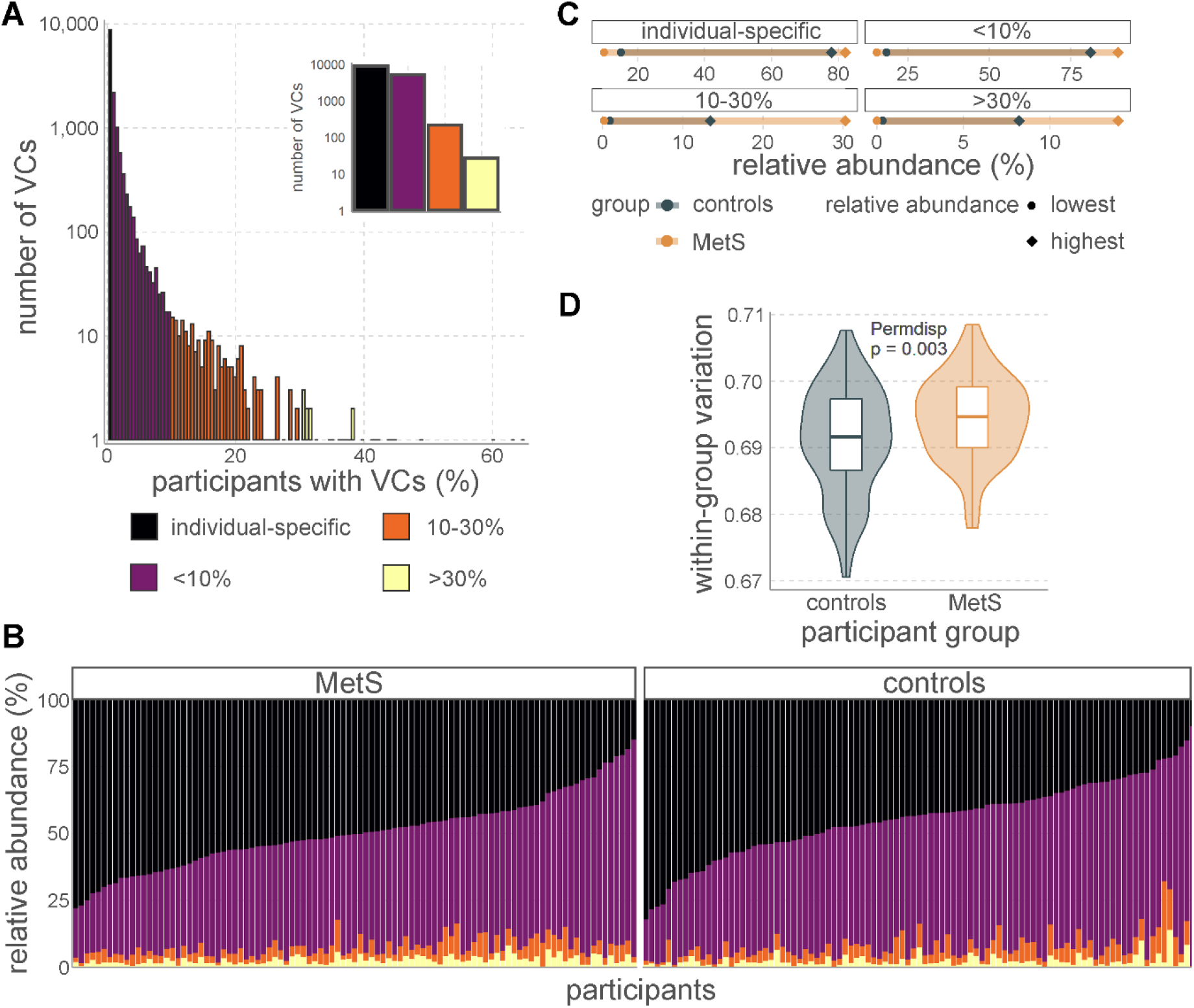
Overview of the phageomes show more variability among MetS participants. **a** Histogram of VCs by number of participants that they are found in shows most VCs are individual-specific. The inset is the same dataset with one bar for each category shown in the legend. **b** Stacked bar charts of community composition show high inter-individual phageome diversity. Color legend is identical to (A). **c** Comparisons show that participants with the highest and lowest relative abundance in each VC category all belong to the MetS group. **d** MetS participants have significantly higher within-group variation, as measured by permdisp on Bray-Curtis dissimilarities. Box plot shows the median, 25^th^, and 75^th^ percentile, with upper and lower whiskers to the 25^th^ percentile minus and the 75^th^ percentile plus 1.5 times the interquartile range.

**Supplementary Figure 2:**
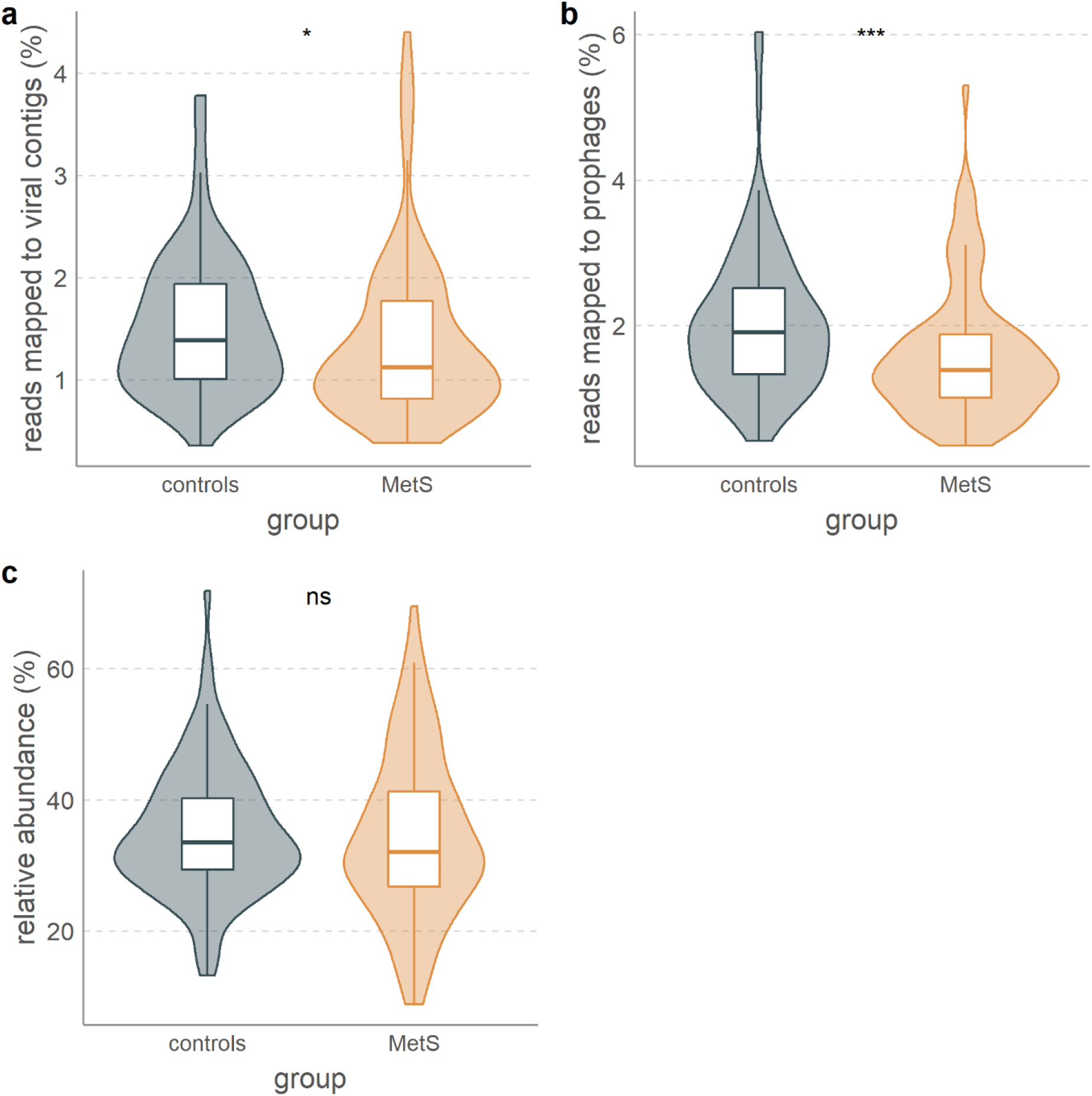
Differences in total phage abundance and potential temperate phage abundance. **a** total phage abundance, as shown by the percentage of reads that map to phage sequences. **b** significantly more reads map to bacterial contigs that contain prophage-like sequences. **c** no significant difference in relative abundance of VCs that carry integrase genes. Stars denote significance according to the Wilcoxon signed rank test. * ≤ 0.05, ** ≤ 0.01, *** ≤ 0.001, **** ≤ 0.0001. Box plots show the median, 25^th^, and 75^th^ percentile, with upper and lower whiskers to the 25^th^ percentile minus and the 75^th^ percentile plus 1.5 times the interquartile range.

**Supplementary Figure 3:**
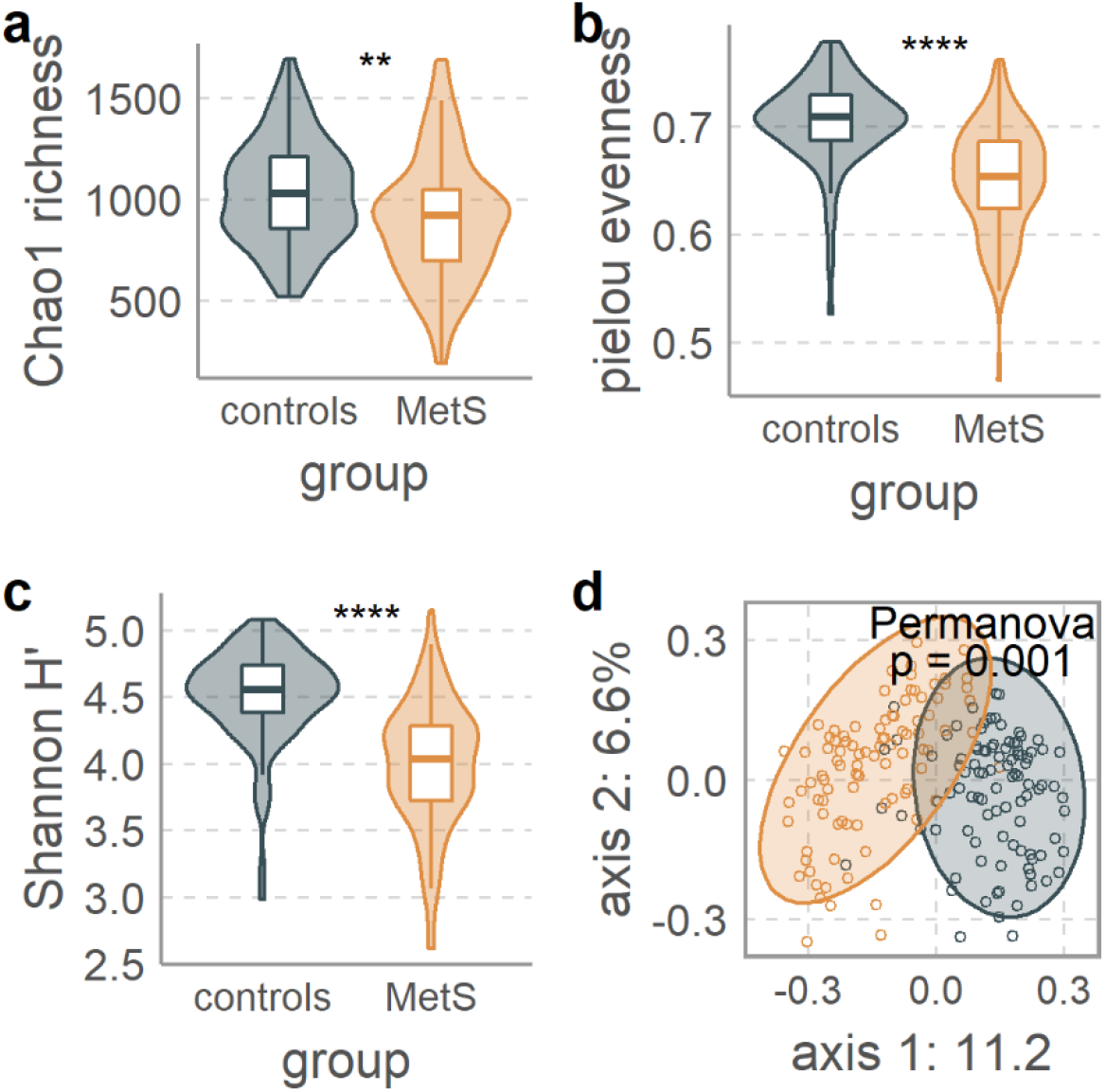
Gut bacterium populations are altered in MetS. **a** MetS-associated decreased bacterial species richness is evidenced by the Chao1 index. **b** decreased bacterial pielou evenness measurements. **c** significantly decreased bacterial α-diversity measured by Shannon diversity. **d** clear separation between bacterial populations of MetS and control participant as shown by β-diversity depicted in a principal coordinates analysis (PCoA) of Bray-Curtis dissimilarities. Permanova test was adjusted for smoking, age, sex, alcohol use, and metformin use. Statistical significance in A-C is according to the Wilcoxon signed rank test, where p-values are denoted as follows: ns not significant, * ≤ 0.05, ** ≤ 0.01, *** ≤ 0.001, **** ≤ 0.0001. Box plots show the median, 25^th^, and 75^th^ percentile, with upper and lower whiskers to the 25^th^ percentile minus and the 75^th^ percentile plus 1.5 times the interquartile range.

**Supplementary Figure 4:**
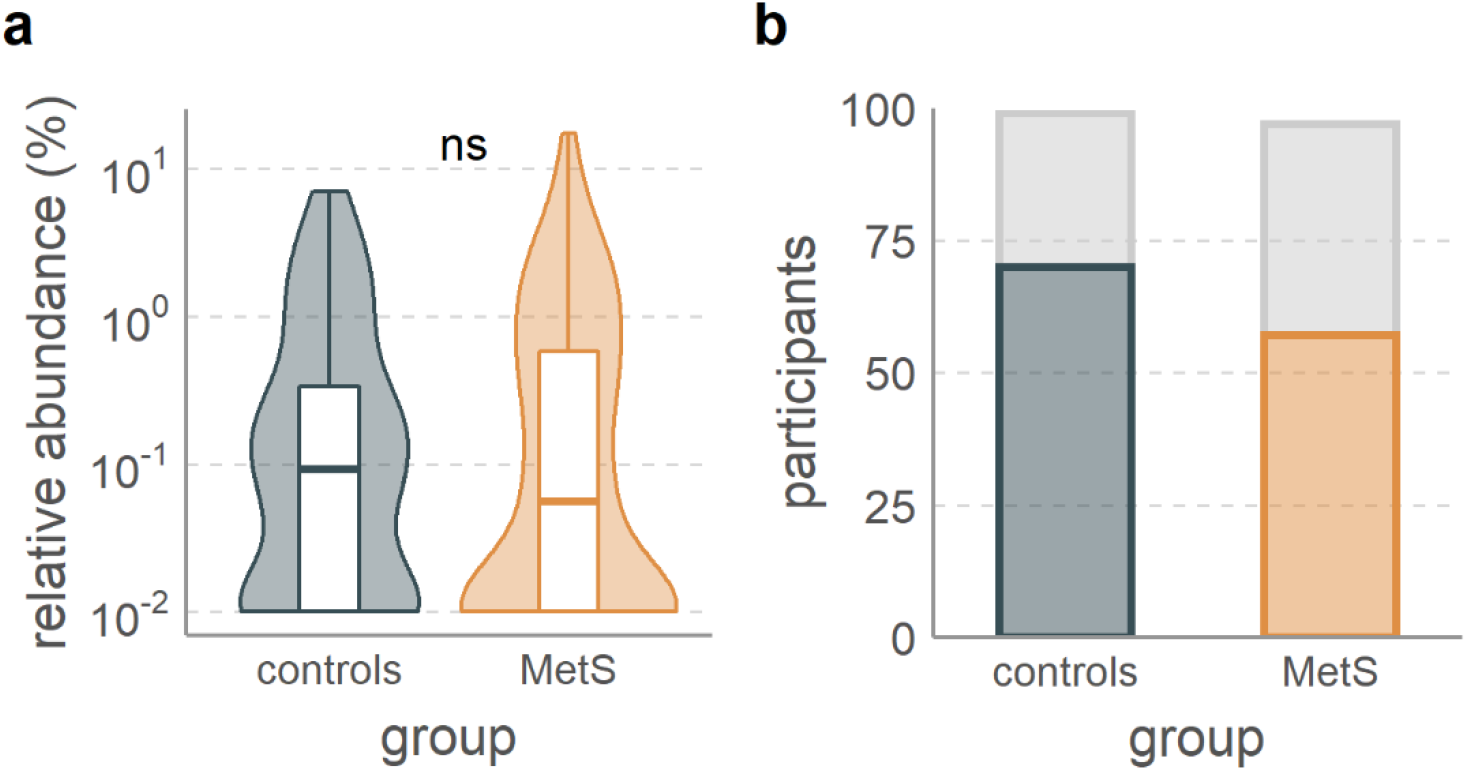
Non-significant differences in crAss-like phage populations. **a** relative abundance of crAss-like phages in controls and MetS. **b** the number of participants in which crAss-like phages were present. Box plots show the median, 25^th^, and 75^th^ percentile, with upper and lower whiskers to the 25^th^ percentile minus and the 75^th^ percentile plus 1.5 times the interquartile range.

**Supplementary Figure 5:**
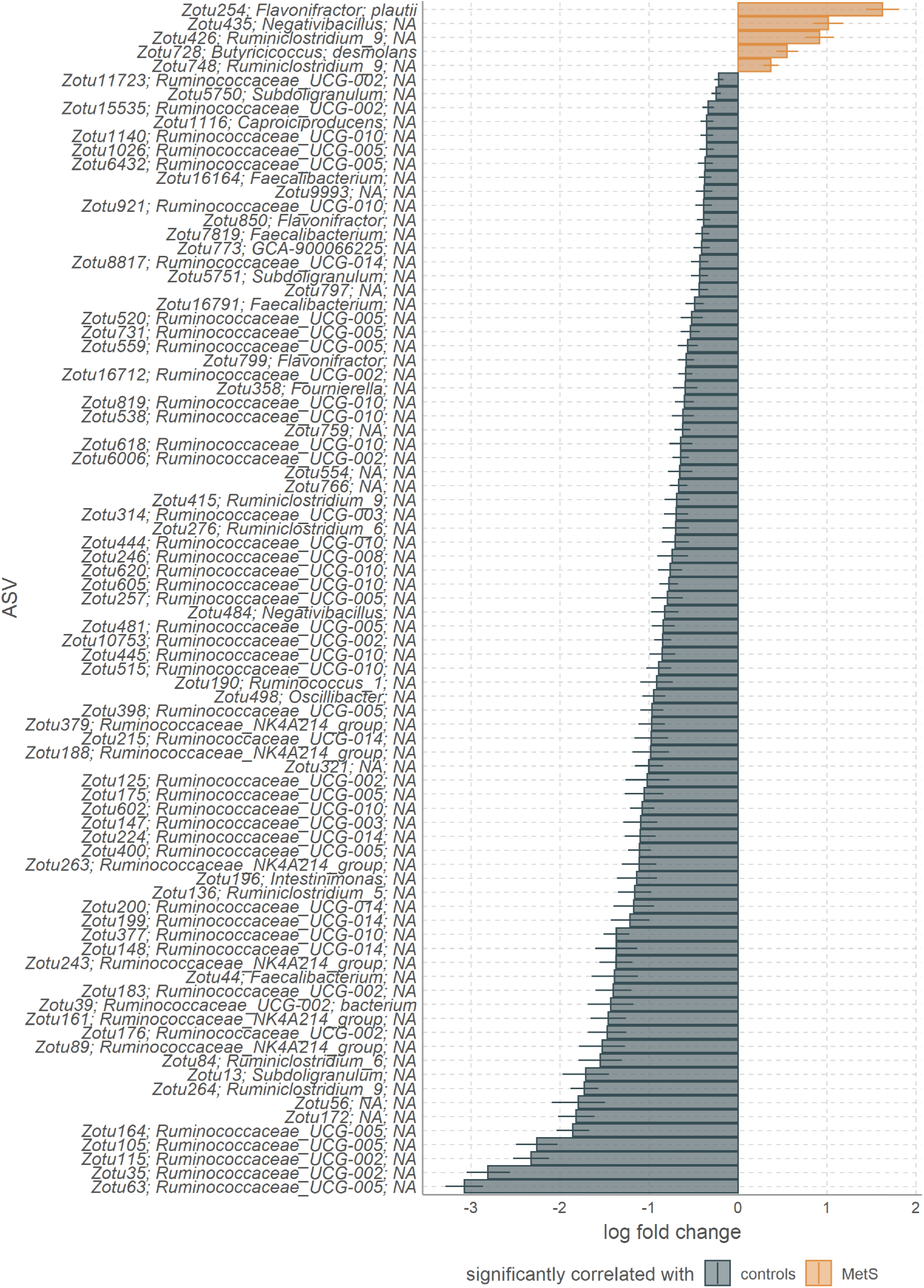
ANCOM-BC analysis results of significantly differentially abundant *Ruminococcaceae* ASVs. Error bars denote the standard error with Holm adjustment for multiple testing.

**Supplementary Figure 6:**
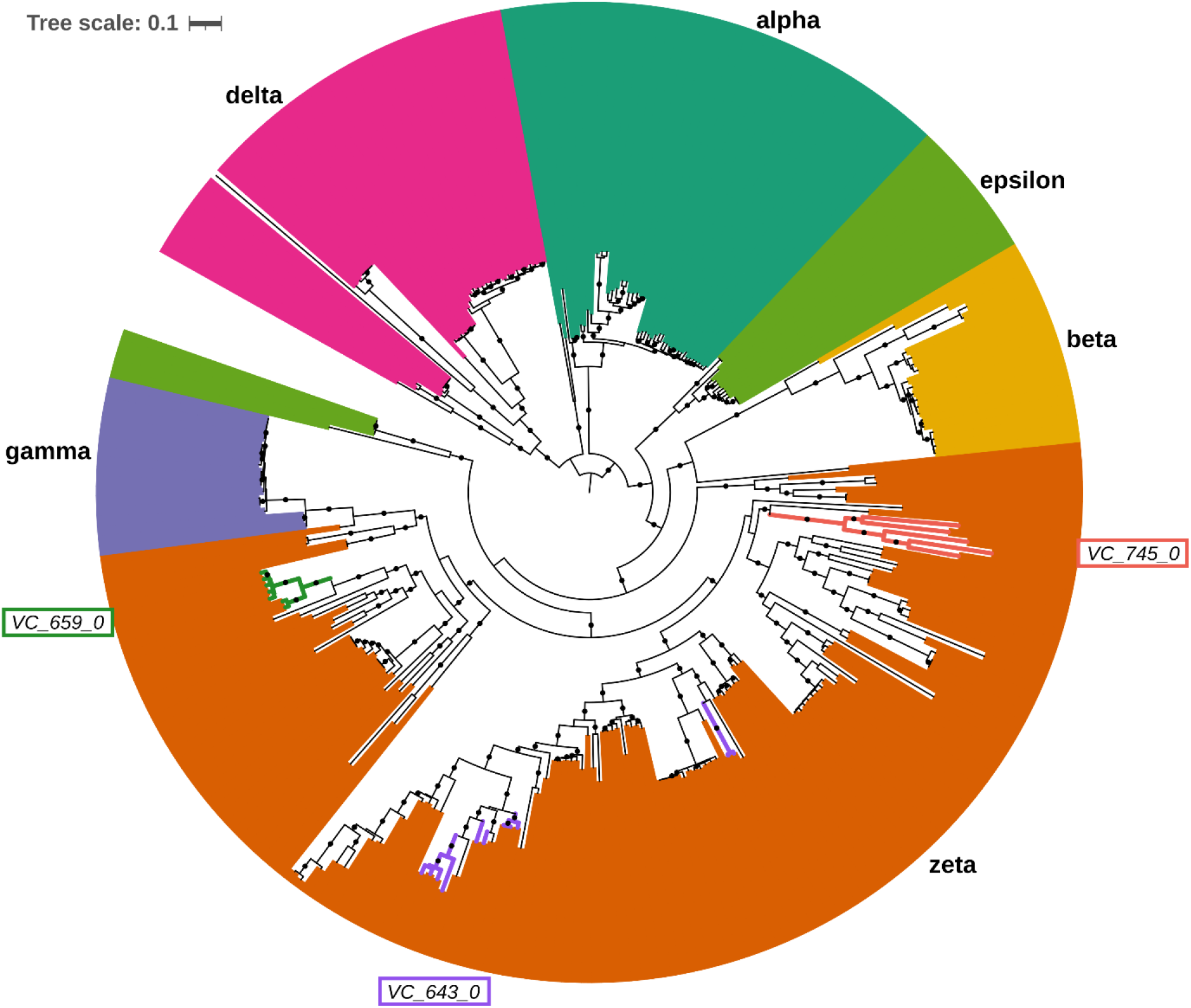
A midpoint-rooted approximate maximum likelihood tree made from the concatenated alignments of the nine universally shared *Candidatus Heliusviridae* genes, with colors denoting the groups. Dots represent bootstrap values of ≥95. Branch colors show contigs that belong to the three *Ca. Heliusviridae* VCs that are significantly differentially abundant in either controls or MetS participants.

**Supplementary Figure 7:**
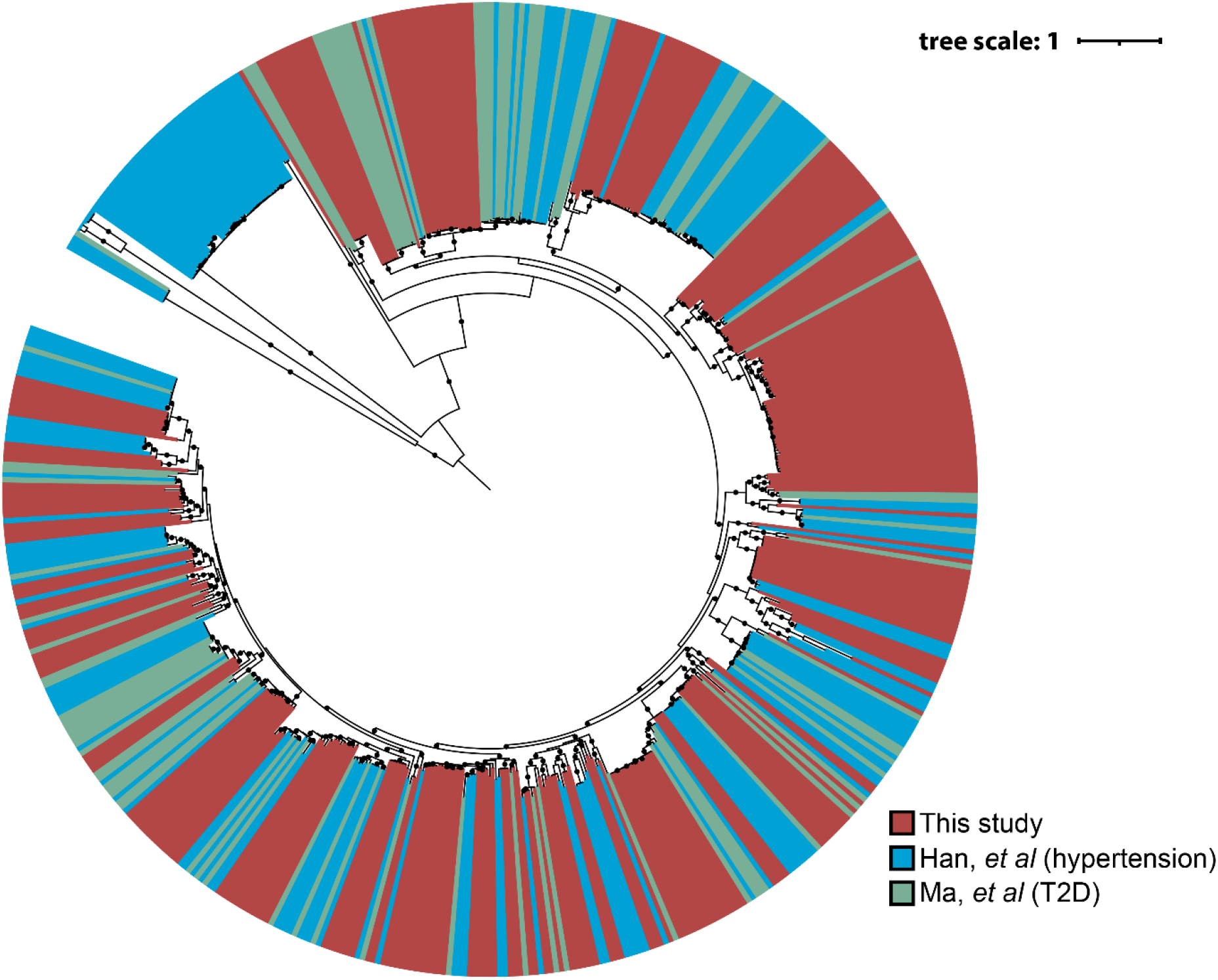
A midpoint-rooted approximate maximum likelihood tree made from the concatenated alignments of the four structural *Candidatus Heliusviridae* genes in contigs from this study and two cohorts in which the phageome was analyzed before, with colors denoting the study. Dots represent bootstrap values of ≥95.

**Supplementary Figure 8:**
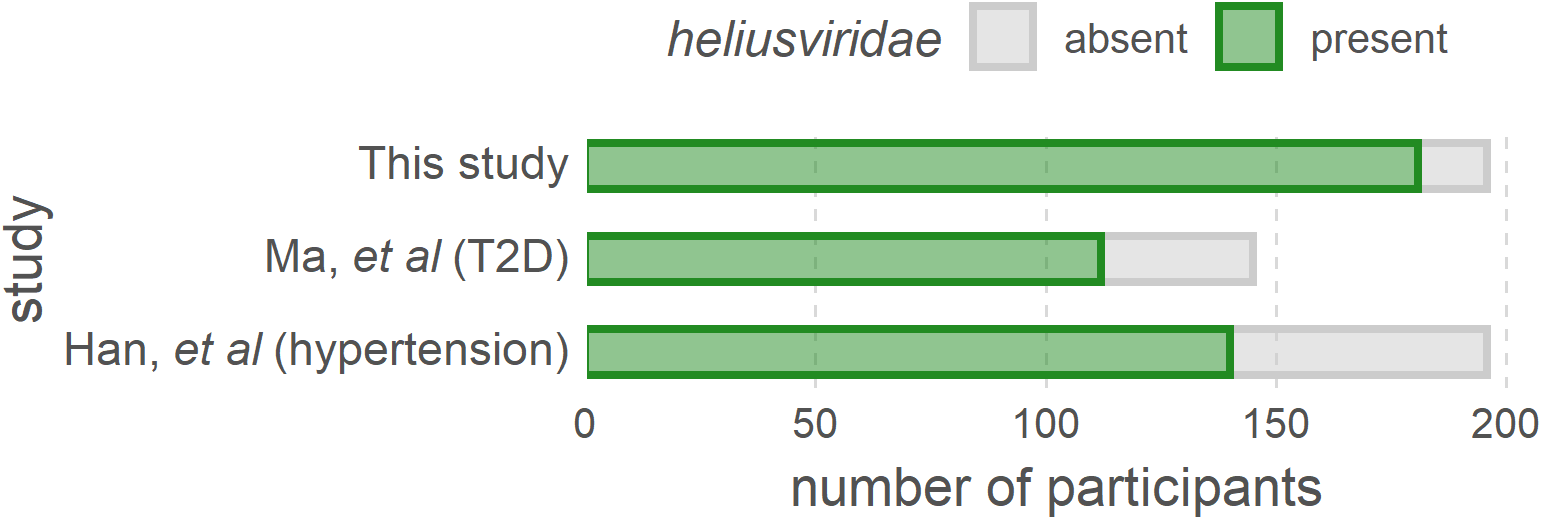
Occurrence of *Candidatus Heliusviridae* in this study and two validation cohorts. To circumvent incomplete assemblies, contigs were identified as *Candidatus Heliusviridae* if they 1) contained the terminase, portal protein, major capsid protein, and clp-proteas, and 2) were located in the same clade as *Candidatus Heliusviridae* from this study in the phylogenetic tree depicted in Supplementary Figure 7.

## Notes

### Competing Interest Statement

The authors have declared no competing interest.

### Summary of Updates

The wording of the text was edited throughout, and we validated the presence of the Ca. Heliusviridae in two independent cohorts, to which effect two supplementary figures were added.

## References

1. Belkaid, Y. & Hand, T. W. Role of the microbiota in immunity and inflammation. Cell 157, 121–141 (2014).

2. Rastelli, M., Cani, P. D. & Knauf, C. The Gut Microbiome Influences Host Endocrine Functions. Endocr. Rev. 40, 1271–1284 (2019).

3. Gurung, M. et al. Role of gut microbiota in type 2 diabetes pathophysiology. EBioMedicine 51, 102590 (2020).

4. Lang, S. & Schnabl, B. Microbiota and Fatty Liver Disease—the Known, the Unknown, and the Future. Cell Host Microbe 28, 233–244 (2020).

5. Frank, D. N. et al. Molecular-phylogenetic characterization of microbial community imbalances in human inflammatory bowel diseases. Proc. Natl. Acad. Sci. 104, 13780–13785 (2007).

6. Clooney, A. G. et al. Whole-Virome Analysis Sheds Light on Viral Dark Matter in Inflammatory Bowel Disease. Cell Host Microbe 26, 764–778.e5 (2019).

7. Norman, J. M. et al. Disease-Specific Alterations in the Enteric Virome in Inflammatory Bowel Disease. Cell 160, 447–460 (2015).

8. Campbell, D. E. et al. Infection with Bacteroides Phage BV01 Alters the Host Transcriptome and Bile Acid Metabolism in a Common Human Gut Microbe. Cell Rep. 32, 108142 (2020).

9. Oh, J.-H. et al. Dietary Fructose and Microbiota-Derived Short-Chain Fatty Acids Promote Bacteriophage Production in the Gut Symbiont Lactobacillus reuteri. Cell Host Microbe 25, 273–284.e6 (2019).

10. Reyes, A. et al. Gut DNA viromes of Malawian twins discordant for severe acute malnutrition. Proc. Natl. Acad. Sci. 112, 11941–11946 (2015).

11. Ma, Y., You, X., Mai, G., Tokuyasu, T. & Liu, C. A human gut phage catalog correlates the gut phageome with type 2 diabetes. Microbiome 6, 1–12 (2018).

12. De Sordi, L., Lourenço, M. & Debarbieux, L. The Battle Within: Interactions of Bacteriophages and Bacteria in the Gastrointestinal Tract. Cell Host Microbe 25, 210–218 (2019).

13. Paez-Espino, D. et al. Uncovering Earth’s virome. Nature 536, 425–430 (2016).

14. Gregory, A. C. et al. The Gut Virome Database Reveals Age-Dependent Patterns of Virome Diversity in the Human Gut. Cell Host Microbe 28, 724–740.e8 (2020).

15. Dutilh, B. E. et al. A highly abundant bacteriophage discovered in the unknown sequences of human faecal metagenomes. Nat. Commun. 5, 4498 (2014).

16. Yutin, N. et al. Discovery of an expansive bacteriophage family that includes the most abundant viruses from the human gut. Nat. Microbiol. 3, 38–46 (2018).

17. O’Neill, S. & O’Driscoll, L. Metabolic syndrome: A closer look at the growing epidemic and its associated pathologies. Obes. Rev. 16, 1–12 (2015).

18. Dabke, K., Hendrick, G. & Devkota, S. The gut microbiome and metabolic syndrome. J. Clin. Invest. 129, 4050–4057 (2019).

19. Mazidi, M., Rezaie, P., Kengne, A. P., Mobarhan, M. G. & Ferns, G. A. Gut microbiome and metabolic syndrome. Diabetes Metab. Syndr. Clin. Res. Rev. 10, S150–S157 (2016).

20. Fujisaka, S. et al. Diet, Genetics, and the Gut Microbiome Drive Dynamic Changes in Plasma Metabolites. Cell Rep. 22, 3072–3086 (2018).

21. Ussar, S. et al. Interactions between gut microbiota, host genetics and diet modulate the predisposition to obesity and metabolic syndrome. Cell Metab. 22, 516–530 (2015).

22. Haro, C. et al. The gut microbial community in metabolic syndrome patients is modified by diet. J. Nutr. Biochem. 27, 27–31 (2016).

23. Shkoporov, A. N. & Hill, C. Bacteriophages of the Human Gut: The “Known Unknown” of the Microbiome. Cell Host Microbe 25, 195–209 (2019).

24. Deschasaux, M. et al. Depicting the composition of gut microbiota in a population with varied ethnic origins but shared geography. Nat. Med. 24, 1526–1531 (2018).

25. Bin Jang, H. et al. Taxonomic assignment of uncultivated prokaryotic virus genomes is enabled by gene-sharing networks. Nat. Biotechnol. 37, 632–639 (2019).

26. Manrique, P. et al. Healthy human gut phageome. Proc. Natl. Acad. Sci. 113, 10400–10405 (2016).

27. Roux, S., Emerson, J. B., Eloe-Fadrosh, E. A. & Sullivan, M. B. Benchmarking viromics: An in silico evaluation of metagenome-enabled estimates of viral community composition and diversity. PeerJ 2017, 1–26 (2017).

28. Camarillo-Guerrero, L. F., Almeida, A., Rangel-Pineros, G., Finn, R. D. & Lawley, T. D. Massive expansion of human gut bacteriophage diversity. Cell 184, 1098–1109.e9 (2021).

29. Lin, H. & Peddada, S. Das. Analysis of compositions of microbiomes with bias correction. Nat. Commun. 11, 1–11 (2020).

30. Aron-Wisnewsky, J. et al. Gut microbiota and human NAFLD: disentangling microbial signatures from metabolic disorders. Nat. Rev. Gastroenterol. Hepatol. 17, 279–297 (2020).

31. Liu, R. et al. Gut microbiome and serum metabolome alterations in obesity and after weight-loss intervention. Nat. Med. 23, 859–868 (2017).

32. Hryckowian, A. J. et al. Bacteroides thetaiotaomicron-Infecting Bacteriophage Isolates Inform Sequence-Based Host Range Predictions. Cell Host Microbe 28, 371–379.e5 (2020).

33. Yutin, N. et al. Analysis of metagenome-assembled viral genomes from the human gut reveals diverse putative CrAss-like phages with unique genomic features. Nat. Commun. 12, 1044 (2021).

34. Nayfach, S. et al. CheckV assesses the quality and completeness of metagenome-assembled viral genomes. Nat. Biotechnol. (2020). doi:10.1038/s41587-020-00774-7

35. Von Meijenfeldt, F. A. B., Arkhipova, K., Cambuy, D. D., Coutinho, F. H. & Dutilh, B. E. Robust taxonomic classification of uncharted microbial sequences and bins with CAT and BAT. Genome Biol. 20, 1–14 (2019).

36. Hedzet, S., Accetto, T. & Rupnik, M. Lytic Bacteroides uniformis bacteriophages exhibiting host tropism congruent with diversity generating retroelement. bioRxiv 2020.10.09.334284 (2020). doi:10.1101/2020.10.09.334284

37. García-López, M. et al. Analysis of 1,000 Type-Strain Genomes Improves Taxonomic Classification of Bacteroidetes. Front. Microbiol. 10, (2019).

38. Maya-Lucas, O. et al. The gut microbiome of Mexican children affected by obesity. Anaerobe 55, 11–23 (2019).

39. Miquel, S. et al. Faecalibacterium prausnitzii and human intestinal health. Curr. Opin. Microbiol. 16, 255–261 (2013).

40. Lavigne, R., Seto, D., Mahadevan, P., Ackermann, H. W. & Kropinski, A. M. Unifying classical and molecular taxonomic classification: analysis of the Podoviridae using BLASTP-based tools. Res. Microbiol. 159, 406–414 (2008).

41. Guerin, E. et al. Biology and Taxonomy of crAss-like Bacteriophages, the Most Abundant Virus in the Human Gut. Cell Host Microbe 24, 653–664.e6 (2018).

42. Han, M., Yang, P., Zhong, C. & Ning, K. The Human Gut Virome in Hypertension. Front. Microbiol. 9, 1–10 (2018).

43. Shkoporov, A. N. et al. The Human Gut Virome Is Highly Diverse, Stable, and Individual Specific. Cell Host Microbe 26, 527–541.e5 (2019).

44. Cornuault, J. K. et al. The enemy from within: a prophage of Roseburia intestinalis systematically turns lytic in the mouse gut, driving bacterial adaptation by CRISPR spacer acquisition. ISME J. 14, 771–787 (2020).

45. Walther, B., Karl, J. P., Booth, S. L. & Boyaval, P. Menaquinones, Bacteria, and the Food Supply: The Relevance of Dairy and Fermented Food Products to Vitamin K Requirements. Adv. Nutr. 4, 463–473 (2013).

46. Moreno-Gallego, J. L. et al. Virome Diversity Correlates with Intestinal Microbiome Diversity in Adult Monozygotic Twins. Cell Host Microbe 25, 261–272.e5 (2019).

47. Zmora, N., Suez, J. & Elinav, E. You are what you eat: diet, health and the gut microbiota. Nat. Rev. Gastroenterol. Hepatol. 16, 35–56 (2019).

48. Falony, G. et al. Population-level analysis of gut microbiome variation. Science (80-.). 352, 560–564 (2016).

49. Zhernakova, A. et al. Population-based metagenomics analysis reveals markers for gut microbiome composition and diversity. Science (80-.). 352, 565–569 (2016).

50. Minot, S. et al. The human gut virome: Inter-individual variation and dynamic response to diet. Genome Res. 21, 1616–1625 (2011).

51. Rodriguez-Valera, F. et al. Explaining microbial population genomics through phage predation. Nat. Rev. Microbiol. 7, 828–836 (2009).

52. Koskella, B. & Brockhurst, M. A. Bacteria–phage coevolution as a driver of ecological and evolutionary processes in microbial communities. FEMS Microbiol. Rev. 38, 916–931 (2014).

53. Silveira, C. B. & Rohwer, F. L. Piggyback-the-Winner in host-associated microbial communities. npj Biofilms Microbiomes 2, 16010 (2016).

54. Fujimoto, K. et al. Metagenome Data on Intestinal Phage-Bacteria Associations Aids the Development of Phage Therapy against Pathobionts. Cell Host Microbe 28, 380–389.e9 (2020).

55. Zaneveld, J. R., McMinds, R. & Vega Thurber, R. Stress and stability: applying the Anna Karenina principle to animal microbiomes. Nat. Microbiol. 2, 17121 (2017).

56. Holmes, I., Harris, K. & Quince, C. Dirichlet Multinomial Mixtures: Generative Models for Microbial Metagenomics. PLoS One 7, e30126 (2012).

57. Lourenço, M. et al. The Spatial Heterogeneity of the Gut Limits Predation and Fosters Coexistence of Bacteria and Bacteriophages. Cell Host Microbe 28, 390–401.e5 (2020).

58. Hatfull, G. F. Dark Matter of the Biosphere: the Amazing World of Bacteriophage Diversity. J. Virol. 89, 8107–8110 (2015).

59. Edwards, R. A., McNair, K., Faust, K., Raes, J. & Dutilh, B. E. Computational approaches to predict bacteriophage-host relationships. FEMS Microbiol. Rev. 40, 258–272 (2016).

60. Burstein, D. et al. Major bacterial lineages are essentially devoid of CRISPR-Cas viral defence systems. Nat. Commun. 7, 10613 (2016).

61. Džunková, M. et al. Defining the human gut host–phage network through single-cell viral tagging. Nat. Microbiol. 4, 2192–2203 (2019).

62. de Jonge, P. A. et al. Adsorption Sequencing as a Rapid Method to Link Environmental Bacteriophages to Hosts. iScience 23, 101439 (2020).

63. Walters, W. A., Xu, Z. & Knight, R. Meta-analyses of human gut microbes associated with obesity and IBD. FEBS Lett. 588, 4223–4233 (2014).

64. Drulis-Kawa, Z., Majkowska-Skrobek, G., Maciejewska, B., Delattre, A.-S. & Lavigne, R. Learning from Bacteriophages - Advantages and Limitations of Phage and Phage-Encoded Protein Applications. Curr. Protein Pept. Sci. 13, 699–722 (2012).

65. Ridaura, V. K. et al. Gut Microbiota from Twins Discordant for Obesity Modulate Metabolism in Mice. Science (80-.). 341, 1241214 (2013).

66. David, L. A. et al. Diet rapidly and reproducibly alters the human gut microbiome. Nature 505, 559–563 (2014).

67. De Filippis, F. et al. Distinct Genetic and Functional Traits of Human Intestinal Prevotella copri Strains Are Associated with Different Habitual Diets. Cell Host Microbe 25, 444–453.e3 (2019).

68. Shkoporov, A. N. et al. ΦCrAss001 represents the most abundant bacteriophage family in the human gut and infects Bacteroides intestinalis. Nat. Commun. 9, 4781 (2018).

69. Koonin, E. V. & Yutin, N. The crAss-like Phage Group: How Metagenomics Reshaped the Human Virome. Trends Microbiol. 28, 349–359 (2020).

70. Edwards, R. A. et al. Global phylogeography and ancient evolution of the widespread human gut virus crAssphage. Nat. Microbiol. 4, 1727–1736 (2019).

71. Garmaeva, S. et al. Studying the gut virome in the metagenomic era: Challenges and perspectives. BMC Biol. 17, 1–14 (2019).

72. Zhao, L. et al. Gut bacteria selectively promoted by dietary fibers alleviate type 2 diabetes. Science (80-.). 359, 1151–1156 (2018).

73. Narita, M. The gut microbiome as a target for prevention of allergic diseases. Japanese J. Allergol. 69, 19–22 (2020).

74. De La Cuesta-Zuluaga, J. et al. Metformin is associated with higher relative abundance of mucin-degrading akkermansia muciniphila and several short-chain fatty acid-producing microbiota in the gut. Diabetes Care 40, 54–62 (2017).

75. Gazitúa, M. C. et al. Potential virus-mediated nitrogen cycling in oxygen-depleted oceanic waters. ISME J. 15, 981–998 (2021).

76. Sharon, I. et al. Photosystem I gene cassettes are present in marine virus genomes. Nature 461, 258–262 (2009).

77. Snijder, M. B. et al. Cohort profile: The Healthy Life in an Urban Setting (HELIUS) study in Amsterdam, the Netherlands. BMJ Open 7, 1–11 (2017).

78. Mobini, R. et al. Metabolic effects of Lactobacillus reuteri DSM 17938 in people with type 2 diabetes: A randomized controlled trial. Diabetes, Obes. Metab. 19, 579–589 (2017).

79. Alberti, K. G. M. M. et al. Harmonizing the metabolic syndrome: A joint interim statement of the international diabetes federation task force on epidemiology and prevention; National heart, lung, and blood institute; American heart association; World heart federation; International. Circulation 120, 1640–1645 (2009).

80. Nurk, S., Meleshko, D., Korobeynikov, A. & Pevzner, P. A. MetaSPAdes: A new versatile metagenomic assembler. Genome Res. 27, 824–834 (2017).

81. Pagès H, Aboyoun P, Gentleman R, D. S. Biostrings: Efficient manipulation of biological strings. (2020).

82. Gregory, A. C. et al. Marine DNA Viral Macro- and Microdiversity from Pole to Pole. Cell 177, 1109–1123.e14 (2019).

83. Roux, S., Enault, F., Hurwitz, B. L. & Sullivan, M. B. VirSorter: mining viral signal from microbial genomic data. PeerJ 3, e985 (2015).

84. Hyatt, D. et al. Prodigal: prokaryotic gene recognition and translation initiation site identification. BMC Bioinformatics 11, 119 (2010).

85. Verhaar, B. J. H. et al. Associations between gut microbiota, faecal short-chain fatty acids, and blood pressure across ethnic groups: the HELIUS study. Eur. Heart J. 41, 4259–4267 (2020).

86. Langmead, B. & Salzberg, S. L. Fast gapped-read alignment with Bowtie 2. Nat. Methods 9, 357–359 (2012).

87. Li, H. et al. The Sequence Alignment/Map format and SAMtools. Bioinformatics 25, 2078–2079 (2009).

88. Quinlan, A. R. BEDTools: The Swiss-Army tool for genome feature analysis. Current Protocols in Bioinformatics 2014, (2014).

89. McMurdie, P. J. & Holmes, S. phyloseq: An R Package for Reproducible Interactive Analysis and Graphics of Microbiome Census Data. PLoS One 8, e61217 (2013).

90. Dixon, P. VEGAN, a package of R functions for community ecology. J. Veg. Sci. 14, 927–930 (2003).

91. Biswas, A., Staals, R. H. J., Morales, S. E., Fineran, P. C. & Brown, C. M. CRISPRDetect: A flexible algorithm to define CRISPR arrays. BMC Genomics 17, 1–14 (2016).

92. Nobrega, F. L., Walinga, H., Dutilh, B. E. & Brouns, S. J. J. J. Prophages are associated with extensive CRISPR–Cas auto-immunity. Nucleic Acids Res. 48, 12074–12084 (2020).

93. Wattam, A. R. et al. PATRIC, the bacterial bioinformatics database and analysis resource. Nucleic Acids Res. 42, 581–591 (2014).

94. Zhang, R. et al. SpacePHARER: Sensitive identification of phages from CRISPR spacers in prokaryotic hosts. bioRxiv 2020.05.15.090266 (2020). doi:10.1101/2020.05.15.090266

95. Pruitt, K. D., Tatusova, T. & Maglott, D. R. NCBI reference sequences (RefSeq): A curated non-redundant sequence database of genomes, transcripts and proteins. Nucleic Acids Res. 35, 61–65 (2007).

96. Gloor, G. B., Macklaim, J. M., Pawlowsky-Glahn, V. & Egozcue, J. J. Microbiome datasets are compositional: And this is not optional. Front. Microbiol. 8, 1–6 (2017).

97. Camacho, C. et al. BLAST+: architecture and applications. BMC Bioinformatics 10, 421 (2009).

98. Sullivan, M. J., Petty, N. K. & Beatson, S. A. Easyfig: A genome comparison visualizer. Bioinformatics 27, 1009–1010 (2011).

99. Seemann, T. Prokka: Rapid prokaryotic genome annotation. Bioinformatics 30, 2068–2069 (2014).

100. Jones, P. et al. InterProScan 5: Genome-scale protein function classification. Bioinformatics 30, 1236–1240 (2014).

101. Johnson, M. et al. NCBI BLAST: a better web interface. Nucleic Acids Res. 36, 5–9 (2008).

102. Sievers, F. & Higgins, D. G. Clustal Omega for making accurate alignments of many protein sequences. Protein Sci. 27, 135–145 (2018).

103. Capella-Gutiérrez, S., Silla-Martínez, J. M. & Gabaldón, T. trimAl: A tool for automated alignment trimming in large-scale phylogenetic analyses. Bioinformatics 25, 1972–1973 (2009).

104. Nguyen, L. T., Schmidt, H. A., Von Haeseler, A. & Minh, B. Q. IQ-TREE: A fast and effective stochastic algorithm for estimating maximum-likelihood phylogenies. Mol. Biol. Evol. 32, 268–274 (2015).

105. Kalyaanamoorthy, S., Minh, B. Q., Wong, T. K. F., von Haeseler, A. & Jermiin, L. S. ModelFinder: fast model selection for accurate phylogenetic estimates. Nat. Methods 14, 587–589 (2017).

106. Hoang, D. T., Chernomor, O., von Haeseler, A., Minh, B. Q. & Vinh, L. S. UFBoot2: Improving the Ultrafast Bootstrap Approximation. Molecular biology and evolution. Mol. Biol. Evol. 35, 518–522 (2018).

107. Zhou, X., Shen, X. X., Hittinger, C. T. & Rokas, A. Evaluating fast maximum likelihood-based phylogenetic programs using empirical phylogenomic data sets. Mol. Biol. Evol. 35, 486–503 (2018).

108. Buchfink, B., Xie, C. & Huson, D. H. Fast and sensitive protein alignment using DIAMOND. Nat. Methods 12, 59–60 (2014).

109. Charrad, M., Ghazzali, N., Boiteau, V. & Niknafs, A. NbClust : An R Package for Determining the Relevant Number of Clusters in a Data Set. J. Stat. Softw. 61, 11744–11750 (2014).

